# Scaling of thermal tolerance with body mass and genome size in ectotherms: A comparison between water-and air-breathers

**DOI:** 10.1101/593293

**Authors:** Félix P. Leiva, Piero Calosi, Wilco C.E.P. Verberk

## Abstract

Global warming appears to favour smaller-bodied organisms, but whether larger species are also more vulnerable to thermal extremes, as suggested for past mass-extinction events, is still an open question. Here, we tested whether interspecific differences in thermal tolerance (heat and cold) of ectotherm organisms are linked to differences in their body mass and genome size (as a proxy for cell size). Since the vulnerability of larger, aquatic taxa to warming has been attributed to the oxygen limitation hypothesis, we also assessed how body mass and genome size modulate thermal tolerance in species with contrasting breathing modes, habitats and life-stages. A database with the upper (CTmax) and lower (CTmin) critical thermal limits and their methodological aspects was assembled comprising more than 500 species of ectotherms. Our results demonstrate that thermal tolerance in ectotherms is dependent on body mass and genome size and these relationships became especially evident in prolonged experimental trials where energy efficiency gains importance. During long-term trials, CTmax was impaired in larger-bodied water-breathers, consistent with a role for oxygen limitation. Variation in CTmin was mostly explained by the combined effects of body mass and genome size and it was enhanced in larger-celled, air-breathing species during long-term trials, consistent with a role for depolarization of cell membranes. Our results highlight the importance of accounting for phylogeny and exposure duration. Especially when considering long-term trials, the observed effects on thermal limits are more in line with the warming-induced reduction in body mass observed during long-term rearing experiments.

## Introduction

The capacity of organisms to take up and transform resources from their environment is a key attribute governing growth, reproduction and subsequently affecting population dynamics, community composition and ecosystem functioning [1,2]. Such capacity seems to be mainly dictated by the species’ body mass [3]. Macroecological and paleoecological data show spatial (e.g. *Bergmann’s rule* [4,5]) and temporal (*Lilliput’s effect* [6]) variation in body mass, which share a common point related to the environmental temperature: at warmer, tropical latitudes and during the past mass extinctions, warming appears to select for smaller-bodied species [5,7–9]. Body size reductions with warming appear to be stronger in aquatic taxa than in terrestrial taxa [5]. In tandem with body size reductions, both aquatic and terrestrial species are shifting their distribution towards cooler habitats and their phenology to earlier and hence, cooler conditions [10,11]. One approach that has been taken to clarify the extent and variation in species redistributions, and to determine which taxonomic groups are potentially more vulnerable to the effects of climate change, is that of comparative studies that analyze thermal tolerance limits (upper and lower) synthesized from the literature [12–15]. These studies also highlight key differences in thermal responses between aquatic and terrestrial taxa, likely related to their breathing mode [16]. The physiological mechanisms underpinning size adjustments and thermal limits are actively debated [17–20], but oxygen limitation has been implicated for both thermal limits [21–23], and size adjustments [24–29] and hypoxia possibly also contributed to mass extinctions [8,30].

By affecting both oxygen demand [31] and the availability of oxygen in water [32,33], warming is hypothesized to result in oxygen limitation, which then causes reductions in thermal limits [22,34] and/or body mass [24,29]. As breathing underwater is more challenging than breathing in air, this oxygen-based mechanism could explain the divergent responses observed in air-and water-breathers [25]. While studies to date hint at a possible size-dependence of thermal limits, no studies have tested this possibility comprehensively. In fact, most studies have focused on one or a few species and although these often find no effect of body mass when included as a covariate in analyses, thermal tolerance limits (heat tolerance rather than cold tolerance) are more frequently reported to decrease rather than increase with increasing body mass [35–38]. In an effort to address this knowledge gap regarding how body mass modulates the response to the temperature in ectotherms, we take advantage of the large body of literature and created a database of upper and lower thermal limits supplemented with biological information of 510 species.

Larger-bodied species may be more susceptible to oxygen limitation because of their lower surface area to volume ratio, which (all else being equal) constrains their capacity to extract oxygen from their environment and deliver it to their metabolizing tissues [24,27,32], or because transport distances increase, which may be especially a problem if these are based on diffusion [28]. If oxygen limitation plays a role in setting thermal limits, one prediction would be that thermal limits vary across organisms with distinct capacities to supply oxygen, including differences between water and air-breathers, or between gas exchange systems across life-stages. As body mass is intimately connected to a suite of other traits, size-dependency of thermal limits may be driven by traits related to body mass, rather than body mass *per se*. For example, relative to the larger adults, smaller life–stages also may experience relatively cooler conditions, especially in temperate and polar regions with a clear seasonality, such that their thermal limits are shifted to lower temperatures, i.e. improved cold tolerance and impaired heat tolerance. Similarly, organisms living in aquatic habitats will experience different thermal regimes than those living on land [15].

Variation in body mass can result from changes in cell number, cell size or a combination of both [39,40], but usually larger-bodied species tend to have larger cells as documented in arthropods [40,41], fish [42] and birds and mammals species [43]. The theory of optimal cell size [44], highlights how differences in cell size have repercussions for oxygen uptake at the cellular level. In the same way, a diversity of cellular physiological functions scales with the cell size [45] and hence, differences in thermal tolerance between animals of different body mass may be mechanistically linked to differences in cell size, rather than body mass. Contrary to the hypothesized influence of oxygen limitation on heat tolerance, the evidence for such an influence on cold tolerance is rather limited [16], and these limits are thought to arise from membrane depolarization and subsequent cell dysfunction due to energy deficits or – in the case of extreme cold tolerance, the freezing of body fluids [46]. Thus, for cold tolerance, a cellular perspective may be more informative, although the correlation between cell size and body mass may result in size-dependency for the CTmin.

In the present study, we use a global database of lower (CTmin) and upper (CTmax) critical thermal limits supplemented with information on other biological traits of ectotherms’ species and their phylogenetic relationships, to investigate whether and how the tolerance to high and low temperatures are modulated by the body mass and genome size (proxy for cell size) across arthropods and vertebrates (amphibians, fish and reptiles) species which distinct breathing-modes, life-stages and habitats. We hypothesize that (1) both CTmax and CTmin will be related to the body mass and genome size of the species, with thermal limits decreasing with increasing body mass (for CTmax) and with increasing genome size (for CTmin); (2) both CTmax and CTmin will differ across breathing modes and a species’ habitat, and such differences will become more pronounced in large-bodied organisms or those with larger genomes; and (3) early life–stages will be more susceptible to heat stress than their adult counterparts, and more resistant to cold stress.

## Materials and methods

### Data search

We created a global database of body mass–related traits (body mass and genome size), life– stage (adult, juvenile and larva) and breathing mode (air, bimodal and water breathers) of aquatic and terrestrial species belonging to four taxonomic groups (amphibians, arthropods, fish and reptiles) for which the critical thermal limits (upper and lower) have been evaluated using dynamic methods (i.e. CTmax or CTmin, *sensu* [47]). The chosen groups comprise taxa for which the determination of body mass was expected to be straightforward. We started by retrieving information from articles on body mass and thermal limits from existing quantitative reviews whose aim has been to explore global patterns of thermal tolerance in ectotherms [12,13,15]. We then added information from recently published references, from January 2015 to October 2018, which were found by using keywords combinations of Boolean terms through ISI Web of Science as follow: (thermal tolerance OR heat tolerance OR cold tolerance OR upper thermal limit OR lower thermal limit OR thermal range OR CTmax OR CTmin) AND (body mass OR body size OR length) AND (amphib* OR arthrop* OR crustacea* OR fish* OR insect* OR reptil*). Searches were limited by research area (ecology, evolutionary biology, biodiversity conservation, environmental sciences, marine freshwater biology, physiology, entomology, zoology, biology, oceanography, fisheries, limnology, environmental studies, behavioural sciences, toxicology, water resources, and multidisciplinary sciences) and research articles. To supplement our searches, we delved into the reference list of each paper to identify additional studies missed in the initial search and if necessary, requesting corresponding authors for additional data not provided in the main text or supplementary information.

### Inclusion criteria

CTmax and CTmin data established by a dynamic (or ramping) method were included in our database, which represents the most common metrics used to assess thermal tolerances in chosen taxa [48]. To account for methodological variation related to differences in starting temperatures and heating/cooling rates across species or studies, we calculated the exposure duration as a single metric that takes into account how long animals are exposed to thermal stress during the heating and cooling trials. After having merged the already published databases with the articles resulted from our search, all duplicates were removed and each article was screened and filtered to build our dataset based only on experimental studies on the basis of three main inclusion criteria: (i) mention of species name belonging to at least one of the four taxa selected (amphibians, arthropods, fish and reptiles), (ii) mention of body mass estimates as mass (wet or dry), width (carapace) or length (carapace, fork, intertegular, snout-vent, standard and total) and (iii) species candidates should be enlisted in the Open Tree of Life (https://ot39.opentreeoflife.org) for subsequent phylogenetic analyses (see “*Data analyses*” section). Despite the restrictive nature of our criteria, just in a few cases, multiple articles reported data on thermal limits for the same species. For this, we prioritized those with the most information available, covering the largest number of entries in our database. Even so, if there were duplicates *per* species, we favored those studies which (i) give both CTmax and CTmin estimations over studies reporting only one thermal limit, (ii) mention the life–stage used during the experiments and (iii) mention methodological information as cooling/heating rates, starting temperatures and geographical coordinates of collection. In the end, all these criteria allowed us to identify 510 species over 174 research articles providing thermal limits and body mass and phylogenetic information (Figure S1, electronic supplementary material). For each species, we compiled taxonomic and biological information (life-stage, habitat, breathing-mode, body mass and genome size), data on the site where a species was collected (geographical coordinates: latitude and longitude, and origin: laboratory or field), methodological information related to the estimation of the thermal limits (starting temperature, heating/cooling rates and acclimation time) and finally, the CTmax and CTmin values.

All body size data collected in units other than mass were transformed using appropriate allometric relationships at the species’ level [49], if it was not possible, we moved up to a higher taxonomic level (e.g. genus or family [50,51]), aiming to have a more representative unit of size for all species in the database, in this case, the body mass in grams (g). As a proxy of cell size, we collected genome size data (in picograms, pg) from the Animal Genome Size Database [52] if it was available. The breathing mode was established on the basis of the species used on each experiment, through “expert judgment” or consulting secondary references if it necessary (e.g. [53]). Bimodal-breathers were classified either as water-breathers (for trials where they relied on under water gas exchange) or air-breathers (for trials where they relied on aerial gas exchange). As most data concerned adults, with only few data for larva and juvenile these two categories were grouped as non-adults. Data from publications where CTmax or CTmin were not reported in the text or tables (i.e. presented only as figures), were extracted using the LibreOffice extension “OOodigitizer v1.2.1”.

### Data analyses

All the results presented in the paper, both in the main text and in the supplementary electronic material were based on linear versions of phylogenetic generalized least squares (PGLSs) models. The correlation structure of these models was given by the potential similarity of species’ traits resulting from the shared evolutionary history and, described by their phylogenetic signal using the Pagel’s lambda (λ) [54]. For this index, a value closer to zero indicates non-phylogenetic signal (phylogenetic independence between species, or a star phylogeny) while a value closer to one indicates that species’ traits evolved randomly through evolutionary time– scales (Brownian phylogeny) [55]. Such information, available as phylogenetic trees, was accessed following [56] and pruned to include only species present in our database. In addition to the estimation of phylogenetic signal in the model residuals, we tested for phylogenetic signal in both in the dependent variables (i.e. the thermal limits) as well as in the independent variables of interest included in the main models following [57] (see electronic supplementary material, Table S11).

Before the main analyses, we first performed preliminary PGLSs in order to determine whether methodological variables influence thermal limits within this dataset and needed to be included in the main analyses. For this, we tested whether the (1) species origin (laboratory or field), or (2) latitude of collection, or (3) acclimation time in the laboratory and the (4) time necessary to reach the CTmax and CTmin affected these thermal limits. The time was calculated after [58,59], as the relation between ramping rate (ΔT, in °C min^-1^) and the starting temperature (T_0_) either for CTmax as: time = [CTmax – T_0_] × ΔT^-1^; and for CTmin trials as: time = [T_0_ – CTmin] × ΔT^-1^. Out of these four methodological variables, only time and/or latitude showed the highest support and also, had significant effects on the thermal limits (for CTmax: latitude and time and for CTmin: only latitude) and these two were subsequently included as covariates in the main models (see electronic supplementary material, Table S1 and Table S2, and Figure S7).

For the main analyses, we fitted PGLSs models each to CTmax and CTmin, first with body mass (log_10_–transformed body mass) as an independent numerical variable, and breathing mode (air and water), life–stage (adult and non-adult) and habitat (aquatic, intertidal and terrestrial) as categorical variables. We also ran models that included all possible interactions of these categorical variables and body mass. In a similar, second set of models, we used genome size (log_10_–transformed genome size) instead of body mass. Since we did not have a reliable estimate of genome size for all 510 taxa, the models using genome size were based on a smaller set of species and hence, model performance cannot be compared directly for those models based on either body mass or genome size.

For each model, we explored the contribution of covariates and phylogeny by fitting models that excluded the effects of phylogeny (i.e. with λ = 0), latitude or exposure duration. Within each variable and covariate combination, we selected the most informative model using a multimodel inference approach by means of the lowest Akaike’s weights (*w*_i_), which provide the relative weight of the evidence towards one of all tested models, and therefore they must add up to 1 [60]. After fitting the models by maximum likelihood, hypothesis testing was performed in models with the highest support using an analysis of deviance with a significance level of *P* ≤ 0.05. All analyses and figures presented in the paper were implemented and generated in R version 3.5.1 [61] using the packages “AICcmodavg” [62], “ape” [63], “nlme” [64], “phytools” [65], “rotl” [56] and “visreg” [66].

## Results

We present results of empirical observations on critical thermal limits for 510 (CTmax) and 232 (CTmin) species (Figures S2a,b, electronic supplementary material). For each species, we also included information on the body mass of the experimental animals (Figure S2c, electronic supplementary material) used during the tests and their phylogenetic relationships (Figures S3-S6, electronic supplementary material). The smallest species (red fire ant *Solenopsis invicta*, 0.0000314 g) is separated from the largest (bonefish *Albula vulpes*, 1235.42 g) by 3.93×10^7^ orders of magnitude (or 7.5 on log_10_-scale). For most of these species, we also included information on their genome size (Figure S2d), and this ranged from 0.14 pg, for the aphid *Aphidius avenae*, to 66.6 pg for the Southern torrent salamander *Rhyacotriton variegatus*. Breathing mode was represented by 225 and 285 species, corresponding to air and water– breathing species, respectively. On the other hand, most data concerned adults (N = 402), while the remaining larvae and juveniles were grouped as non-adults (N = 108). In terms of habitat, the majority of species were aquatic (316 species), or terrestrial (181 species), with only a low species being intertidal (13 species).

Both CTmax and CTmin showed a clear phylogenetic signal (Table S11, electronic supplementary material), indicating that thermal tolerance among the studied species has been largely conserved across evolutionary lineages. A comparison between phylogenetic generalized least squares (PGLS) models under a Brownian mode of evolution (λ = 1) and non-phylogenetic models (λ = 0) showed, in most of cases, that accounting for phylogenetic relationships among the studied species improved the model fit both for CTmax (Table 1 and Table 3) and CTmin (Table 2 and Table 4). The two covariates (i.e. absolute latitude and exposure duration) were always included in the best-supported model, indicating their importance in explaining variation in thermal tolerance. For all CTmax models, greater support and the lowest AICc were observed when phylogeny was taken into account (λ = 1). By contrast, for CTmin, accounting for the shared evolutionary history of species was less important for those models that already included body mass as an explanatory variable, possibly because body mass is strongly phylogenetically structured and may, therefore, obviate the need to include phylogeny (Table 2, model 5 to model 10).

**Table 1.** Results for phylogenetic generalized least squares (PGLSs) models to explain variation in ectotherms’ CTmax (N = 510 species) as a function of log_10_–transformed body mass, breathing mode (air and water), life-stage (adult and non-adult), habitat (aquatic, intertidal and terrestrial) and their interactions. All models were assessed using exposure duration (Time) and/or absolute latitude (Lat) of the animal collection as covariates. The number of parameters (*k*), corrected Akaike’s information criterion (AICc), the difference in AICc respect to the model with highest support (ΔAICc) and the Akaike’s weight (*w*_i_) are mentioned for each model. Pagel’s lambda (λ) denotes correlation structure used (λ = 0, star phylogeny and λ = 1, Brownian phylogeny).

| Models | | *k* | AICc | ΔAICc | *w<sub>i</sub>* |
| --- | --- | --- | --- | --- | --- |
| 0. Covariates | $\lambda = 1 + \text{Lat} + \text{Time}$ | 4 | 3013.58 | 54.43 | 0.00 |
| | $\lambda = 0 + \text{Lat} + \text{Time}$ | 4 | 3306.34 | 347.19 | 0.00 |
| | $\lambda = 1 + \text{Lat}$ | 3 | 3040.98 | 81.83 | 0.00 |
| | $\lambda = 1 + \text{Time}$ | 3 | 3066.19 | 107.04 | 0.00 |
| 1. Body mass | $\lambda = 1 + \text{Lat} + \text{Time}$ | 5 | 3014.70 | 55.55 | 0.00 |
| | $\lambda = 0 + \text{Lat} + \text{Time}$ | 5 | 3200.72 | 241.57 | 0.00 |
| | $\lambda = 1 + \text{Lat}$ | 4 | 3042.19 | 83.04 | 0.00 |
| | $\lambda = 1 + \text{Time}$ | 4 | 3066.02 | 106.87 | 0.00 |
| 2. Breathing mode | $\lambda = 1 + \text{Lat} + \text{Time}$ | 5 | 2984.49 | 25.34 | 0.00 |
| | $\lambda = 0 + \text{Lat} + \text{Time}$ | 5 | 3232.59 | 273.43 | 0.00 |
| | $\lambda = 1 + \text{Lat}$ | 4 | 3005.53 | 46.38 | 0.00 |
| | $\lambda = 1 + \text{Time}$ | 4 | 3031.19 | 72.03 | 0.00 |
| 3. Life-stage | $\lambda = 1 + \text{Lat} + \text{Time}$ | 5 | 3015.52 | 56.37 | 0.00 |
| | $\lambda = 0 + \text{Lat} + \text{Time}$ | 5 | 3307.18 | 348.02 | 0.00 |
| | $\lambda = 1 + \text{Lat}$ | 4 | 3039.31 | 80.16 | 0.00 |
| | $\lambda = 1 + \text{Time}$ | 4 | 3067.71 | 108.56 | 0.00 |
| 4. Habitat | $\lambda = 1 + \text{Lat} + \text{Time}$ | 6 | 3007.04 | 47.89 | 0.00 |
| | $\lambda = 0 + \text{Lat} + \text{Time}$ | 6 | 3246.43 | 287.27 | 0.00 |
| | $\lambda = 1 + \text{Lat}$ | 5 | 3032.57 | 73.42 | 0.00 |
| | $\lambda = 1 + \text{Time}$ | 5 | 3063.24 | 104.09 | 0.00 |
| 5. Body mass × Breathing mode | $\lambda = 1 + \text{Lat} + \text{Time}$ | 7 | 2980.31 | 21.16 | 0.00 |
| | $\lambda = 0 + \text{Lat} + \text{Time}$ | 7 | 3170.76 | 211.61 | 0.00 |
| | $\lambda = 1 + \text{Lat}$ | 6 | 3002.48 | 43.33 | 0.00 |
| | $\lambda = 1 + \text{Time}$ | 6 | 3030.55 | 71.40 | 0.00 |
| 6. Body mass × Life-stage | $\lambda = 1 + \text{Lat} + \text{Time}$ | 7 | 3004.43 | 45.27 | 0.00 |
| | $\lambda = 0 + \text{Lat} + \text{Time}$ | 7 | 3190.02 | 230.87 | 0.00 |
| | $\lambda = 1 + \text{Lat}$ | 6 | 3031.97 | 72.82 | 0.00 |
| | $\lambda = 1 + \text{Time}$ | 6 | 3056.17 | 97.02 | 0.00 |
| 7. Body mass × Habitat | $\lambda = 1 + \text{Lat} + \text{Time}$ | 9 | 3003.94 | 44.79 | 0.00 |
| | $\lambda = 0 + \text{Lat} + \text{Time}$ | 9 | 3167.62 | 208.47 | 0.00 |
| | $\lambda = 1 + \text{Lat}$ | 8 | 3027.88 | 68.73 | 0.00 |
| | $\lambda = 1 + \text{Time}$ | 8 | 3052.40 | 93.25 | 0.00 |
| **8. Body mass × Breathing mode × Time** | **$\lambda = 1 + \text{Lat}$** | **10** | **2959.15** | **0.00** | **1.00** |
|  | **$\lambda = 0 + \text{Lat}$** | **10** | **3148.75** | **189.60** | **0.00** |
| 9. Body mass × Life-stage × Time | $\lambda = 1 + \text{Lat}$ | 10 | 2991.95 | 32.80 | 0.00 |
| | $\lambda = 0 + \text{Lat}$ | 10 | 3189.60 | 230.44 | 0.00 |
| 10. Body mass × Habitat × Time | $\lambda = 1 + \text{Lat}$ | 14 | 2976.66 | 17.51 | 0.00 |
| | $\lambda = 0 + \text{Lat}$ | 14 | 3174.65 | 215.49 | 0.00 |

**Table 2.** Results for phylogenetic generalized least squares (PGLSs) models to explain variation in ectotherms’ CTmin (N = 232 species) as a function of log_10_–transformed body mass, breathing mode (air and water), life-stage (adult and non-adult), habitat (aquatic, intertidal and terrestrial), exposure duration (Time) and their interactions. All models were assessed using absolute latitude (Lat) of the animal collection as a covariate. The number of parameters (*k*), corrected Akaike’s information criterion (AICc), the difference in AICc respect to the model with highest support (ΔAICc) and Akaike’s weight (*w*_i_) are mentioned for each model. Pagel’s lambda (λ) denotes correlation structure used (λ = 0, star phylogeny and λ = 1, Brownian phylogeny).

| Models | | *k* | AICc | ΔAICc | $w_i$ |
| --- | --- | --- | --- | --- | --- |
| 0. Covariates | $\lambda = 1 + \text{Lat}$ | 3 | 1304.72 | 69.18 | 0.00 |
| | $\lambda = 0 + \text{Lat}$ | 3 | 1399.46 | 163.92 | 0.00 |
| 1. Body mass | $\lambda = 1 + \text{Lat}$ | 4 | 1306.73 | 71.19 | 0.00 |
| | $\lambda = 0 + \text{Lat}$ | 4 | 1289.78 | 54.24 | 0.00 |
| 2. Breathing Mode | $\lambda = 1 + \text{Lat}$ | 4 | 1305.39 | 69.85 | 0.00 |
| | $\lambda = 0 + \text{Lat}$ | 4 | 1384.55 | 149.01 | 0.00 |
| 3. Life-stage | $\lambda = 1 + \text{Lat}$ | 4 | 1306.57 | 71.03 | 0.00 |
| | $\lambda = 0 + \text{Lat}$ | 4 | 1398.91 | 163.37 | 0.00 |
| 4. Habitat | $\lambda = 1 + \text{Lat}$ | 5 | 1292.00 | 56.46 | 0.00 |
| | $\lambda = 0 + \text{Lat}$ | 5 | 1396.35 | 160.81 | 0.00 |
| 5. Body mass × Breathing mode | $\lambda = 1 + \text{Lat}$ | 6 | 1307.00 | 71.46 | 0.00 |
| | $\lambda = 0 + \text{Lat}$ | 6 | 1273.49 | 37.95 | 0.00 |
| 6. Body mass × Life-stage | $\lambda = 1 + \text{Lat}$ | 6 | 1306.84 | 71.30 | 0.00 |
| | $\lambda = 0 + \text{Lat}$ | 6 | 1285.09 | 49.55 | 0.00 |
| 7. Body mass × Habitat | $\lambda = 1 + \text{Lat}$ | 8 | 1297.78 | 62.24 | 0.00 |
| | $\lambda = 0 + \text{Lat}$ | 8 | 1274.84 | 39.30 | 0.00 |
| **8. Body mass × Breathing mode × Time** | $\lambda = 1 + \text{Lat}$ | 10 | 1288.19 | 52.65 | 0.00 |
|  | **$\lambda = 0 + \text{Lat}$** | **10** | **1235.54** | **0.00** | **1.00** |
| 9. Body mass × Life-stage × Time | $\lambda = 1 + \text{Lat}$ | 10 | 1290.22 | 54.68 | 0.00 |
| | $\lambda = 0 + \text{Lat}$ | 10 | 1249.66 | 14.12 | 0.00 |
| 10. Body mass × Habitat × Time | $\lambda = 1 + \text{Lat}$ | 14 | 1269.39 | 33.85 | 0.00 |
| | $\lambda = 0 + \text{Lat}$ | 14 | 1264.29 | 28.75 | 0.00 |

**Table 3.** Results for phylogenetic generalized least squares (PGLSs) models to explain variation in ectotherms’ CTmax (N= 433 species) as a function of log_10_–transformed genome size, breathing mode (air and water), life-stage (adult, non-adult), habitat (aquatic, intertidal and terrestrial) and their interactions. All models were assessed using exposure duration (Time) and/or absolute latitude (Lat) of the animal collection as covariates. The number of parameters (*k*), corrected Akaike’s information criterion (AICc), the difference in AICc respect to the model with highest support (ΔAICc) and Akaike’s weight (*w*_i_) are mentioned for each model. Pagel’s lambda (λ) denotes correlation structure used (λ = 0, star phylogeny and λ = 1, Brownian phylogeny).

| **Models** |  | ***k*** | **AICc** | **ΔAICc** | ***w<sub>i</sub>*** |
| --- | --- | --- | --- | --- | --- |
| 0. Covariates | $\lambda = 1 + \text{Lat} + \text{Time}$ | 4 | 2505.59 | 3.24 | 0.07 |
| | $\lambda = 0 + \text{Lat} + \text{Time}$ | 4 | 2759.27 | 256.92 | 0.00 |
| | $\lambda = 1 + \text{Lat}$ | 3 | 2521.96 | 19.60 | 0.00 |
| | $\lambda = 1 + \text{Time}$ | 3 | 2555.30 | 52.94 | 0.00 |
| 1. Genome size | $\lambda = 1 + \text{Lat} + \text{Time}$ | 5 | 2506.18 | 3.82 | 0.05 |
| | $\lambda = 0 + \text{Lat} + \text{Time}$ | 5 | 2731.05 | 228.69 | 0.00 |
| | $\lambda = 1 + \text{Lat}$ | 4 | 2521.94 | 19.58 | 0.00 |
| | $\lambda = 1 + \text{Time}$ | 4 | 2554.25 | 51.89 | 0.00 |
| **2. Breathing mode** | **$\lambda = 1 + \text{Lat} + \text{Time}$** | **5** | **2504.30** | **1.95** | **0.13** |
| | $\lambda = 0 + \text{Lat} + \text{Time}$ | 5 | 2644.82 | 142.46 | 0.00 |
| | $\lambda = 1 + \text{Lat}$ | 4 | 2519.78 | 17.43 | 0.00 |
| | $\lambda = 1 + \text{Time}$ | 4 | 2551.15 | 48.79 | 0.00 |
| **3. Life-stage** | **$\lambda = 1 + \text{Lat} + \text{Time}$** | **5** | **2503.91** | **1.55** | **0.16** |
| | $\lambda = 0 + \text{Lat} + \text{Time}$ | 5 | 2758.50 | 256.14 | 0.00 |
| | $\lambda = 1 + \text{Lat}$ | 4 | 2513.96 | 11.60 | 0.00 |
| | $\lambda = 1 + \text{Time}$ | 4 | 2551.56 | 49.20 | 0.00 |
| 4. Habitat | $\lambda = 1 + \text{Lat} + \text{Time}$ | 6 | 2509.48 | 7.12 | 0.01 |
| | $\lambda = 0 + \text{Lat} + \text{Time}$ | 6 | 2718.37 | 216.01 | 0.00 |
| | $\lambda = 1 + \text{Lat}$ | 5 | 2525.79 | 23.43 | 0.00 |
| | $\lambda = 1 + \text{Time}$ | 5 | 2558.64 | 56.28 | 0.00 |
| 5. Genome size × Breathing mode | $\lambda = 1 + \text{Lat} + \text{Time}$ | 7 | 2506.43 | 4.07 | 0.05 |
| | $\lambda = 0 + \text{Lat} + \text{Time}$ | 7 | 2599.70 | 97.34 | 0.00 |
| | $\lambda = 1 + \text{Lat}$ | 6 | 2521.26 | 18.90 | 0.00 |
| | $\lambda = 1 + \text{Time}$ | 6 | 2551.27 | 48.91 | 0.00 |
| 6. Genome size × Life-stage | $\lambda = 1 + \text{Lat} + \text{Time}$ | 7 | 2506.65 | 4.30 | 0.04 |
| | $\lambda = 0 + \text{Lat} + \text{Time}$ | 7 | 2715.88 | 213.52 | 0.00 |
| | $\lambda = 1 + \text{Lat}$ | 6 | 2515.68 | 13.32 | 0.00 |
| | $\lambda = 1 + \text{Time}$ | 6 | 2552.49 | 50.13 | 0.00 |
| 7. Genome size × Habitat | $\lambda = 1 + \text{Lat} + \text{Time}$ | 9 | 2511.67 | 9.31 | 0.00 |
| | $\lambda = 0 + \text{Lat} + \text{Time}$ | 9 | 2669.46 | 167.10 | 0.00 |
| | $\lambda = 1 + \text{Lat}$ | 8 | 2527.59 | 25.23 | 0.00 |
| | $\lambda = 1 + \text{Time}$ | 8 | 2553.99 | 51.64 | 0.00 |
| **8. Genome size × Breathing mode × Time** | **$\lambda = 1 + \text{Lat}$** | **10** | **2502.36** | **0.00** | **0.36** |
| | $\lambda = 0 + \text{Lat}$ | 10 | 2590.26 | 87.90 | 0.00 |
| 9. Genome size × Life-stage × Time | $\lambda = 1 + \text{Lat}$ | 10 | 2510.08 | 7.72 | 0.01 |
| | $\lambda = 0 + \text{Lat}$ | 10 | 2718.49 | 216.13 | 0.00 |
| 10. Genome size × Habitat × Time | $\lambda = 1 + \text{Lat}$ | 14 | 2504.72 | 2.36 | 0.11 |
| | $\lambda = 0 + \text{Lat}$ | 14 | 2669.50 | 167.14 | 0.00 |

**Table 4.** Results for phylogenetic generalized least squares (PGLSs) models to explain variation in ectotherms’ CTmin (N = 190 species) as a function of log_10_–transformed genome size, breathing mode (air and water), life-stage (adult and non-adult), habitat (aquatic, intertidal and terrestrial) and their interactions. All models were assessed using the absolute latitude (Lat) of the animal collection as a covariate. The number of parameters (*k*), corrected Akaike’s information criterion (AICc), the difference in AICc respect to the model with highest support (ΔAICc) and Akaike’s weight (*w*_i_) are mentioned for each model. Pagel’s lambda (λ) denotes correlation structure used (λ = 0, star phylogeny and λ = 1, Brownian phylogeny).

| Models | | *k* | AICc | ΔAICc | *w<sub>i</sub>* |
| --- | --- | --- | --- | --- | --- |
| 0. Covariates | $\lambda = 1 + \text{Lat}$ | 3 | 1076.21 | 37.66 | 0.00 |
| | $\lambda = 0 + \text{Lat}$ | 3 | 1157.44 | 118.88 | 0.00 |
| 1. Genome size | $\lambda = 1 + \text{Lat}$ | 4 | 1074.03 | 35.47 | 0.00 |
| | $\lambda = 0 + \text{Lat}$ | 4 | 1159.53 | 120.97 | 0.00 |
| 2. Breathing Mode | $\lambda = 1 + \text{Lat}$ | 4 | 1076.83 | 38.27 | 0.00 |
| | $\lambda = 0 + \text{Lat}$ | 4 | 1150.57 | 112.01 | 0.00 |
| 3. Life-stage | $\lambda = 1 + \text{Lat}$ | 4 | 1077.85 | 39.29 | 0.00 |
| | $\lambda = 0 + \text{Lat}$ | 4 | 1158.68 | 120.13 | 0.00 |
| 4. Habitat | $\lambda = 1 + \text{Lat}$ | 5 | 1063.79 | 25.23 | 0.00 |
| | $\lambda = 0 + \text{Lat}$ | 5 | 1148.50 | 109.94 | 0.00 |
| 5. Genome size × Breathing Mode | $\lambda = 1 + \text{Lat}$ | 6 | 1070.58 | 32.02 | 0.00 |
| | $\lambda = 0 + \text{Lat}$ | 6 | 1144.42 | 105.87 | 0.00 |
| 6. Genome size × Life-stage | $\lambda = 1 + \text{Lat}$ | 6 | 1075.65 | 37.09 | 0.00 |
| | $\lambda = 0 + \text{Lat}$ | 6 | 1154.62 | 116.06 | 0.00 |
| 7. Genome size × Habitat | $\lambda = 1 + \text{Lat}$ | 8 | 1046.08 | 7.53 | 0.02 |
| | $\lambda = 0 + \text{Lat}$ | 8 | 1153.09 | 114.53 | 0.00 |
| 8. Genome size × Breathing mode × Time | $\lambda = 1 + \text{Lat}$ | 10 | 1047.97 | 9.42 | 0.01 |
| | $\lambda = 0 + \text{Lat}$ | 10 | 1136.59 | 98.03 | 0.00 |
| 9. Genome size × Life-stage × Time | $\lambda = 1 + \text{Lat}$ | 10 | 1045.99 | 7.43 | 0.02 |
| | $\lambda = 0 + \text{Lat}$ | 10 | 1156.36 | 117.80 | 0.00 |
| **10. Genome size × Habitat × Time** | **$\lambda = 1 + \text{Lat}$** | **14** | **1038.56** | **0.00** | **0.95** |
| | $\lambda = 0 + \text{Lat}$ | 14 | 1153.79 | 115.23 | 0.00 |

Modelled effects of body mass and genome size on both thermal limits differed according to whether the model included phylogeny or not. For CTmax, a negative relationships with body mass was most apparent in the model that did not include phylogeny (λ = 0), likely because both extreme values for CTmax and body mass were phylogenetically clustered (Table 1 and Table 3; Figure 1a,b; Figures S3-S4, electronic supplementary material). Effects of both body mass and genome size on CTmax differed with breathing mode and exposure duration (Tables S3 and S5; see below). For CTmin, the best-supported models indicated that cold tolerance declined (i.e. higher CTmin values) with increasing body mass (Table 2 and Figure 1c) and with increasing genome size (Table 4 and Figure 1d). Effects of body mass on CTmin differed with breathing mode and exposure duration (Table S4), whereas those of genome size differed with habitat and exposure duration (Table S6).

**Figure 1.**
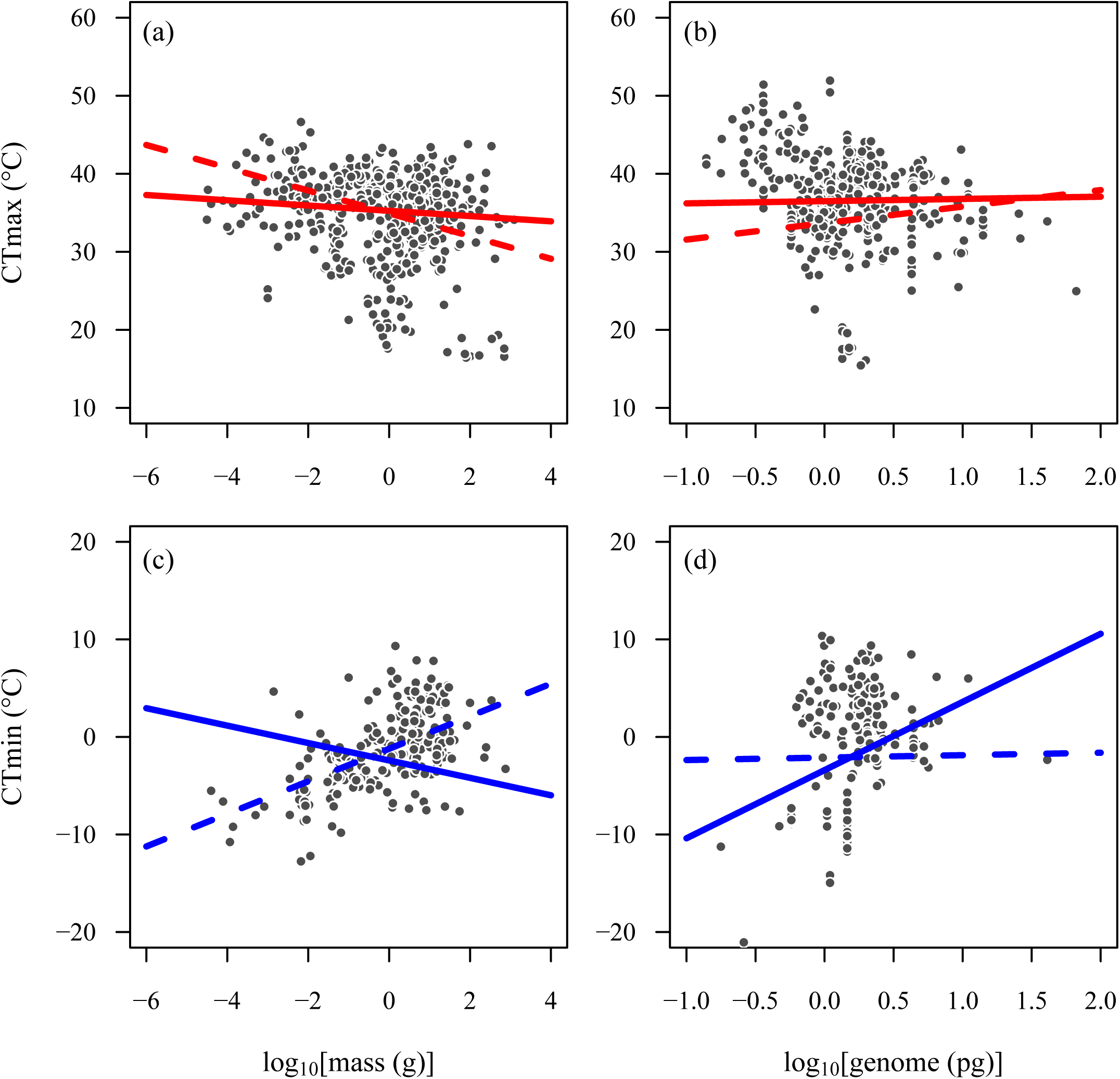
Partial residuals plots showing the predicted effects of log_10_–transformed body mass and log_10_–transformed genome size in ectotherms’ CTmax (top, red) and CTmin (bottom, blue). CTmax (a, b) and CTmin plots (c, d) were based on the model with the highest support shown in Tables 1–4. Solid lines indicate predictions of models that included all covariates (latitude, time) and phylogenetic relationships, whereas dashed lines indicate predictions of models that included all covariates, but do not account for phylogenetic relationships (λ = 0). For details on model estimates and significance, see electronic supplementary material Tables S3-S6.

Consistent with the expectation that both CTmax and CTmin differ in species with different breathing modes and across habitats, our results indicate a generally lower tolerance for water-breathers compared to air-breathers, suggesting that water-breathers are more vulnerable to both heat and cold (Figure 2a,d). Contrary to our expectation, we found no differences in thermal limits between different life-stages (Figure 2b,e). Intertidal species were shown to be more tolerant to the effects of cold (Figure 2f). However, these results should be interpreted with caution in light of low representation of intertidal species in our analyses (5 species for CTmin). Also, this difference for intertidal species was no longer present when phylogenetic relationships were not accounted for (Figure S8, electronic supplementary material). Although breathing mode and habitat strongly covaried (most aquatic species are water-breathers and most terrestrial species are air-breathers), variation in CTmax was best explained by models based on breathing mode (Table 1, model 8), not habitat (Table 1, model 10). Variation in CTmin was best explained by models based on breathing mode (when including body mass; Table 2) and habitat (when including genome size; Table 4). Cold tolerance declined (i.e. higher CTmin values) with increasing body mass (Figure 1c).

**Figure 2.**
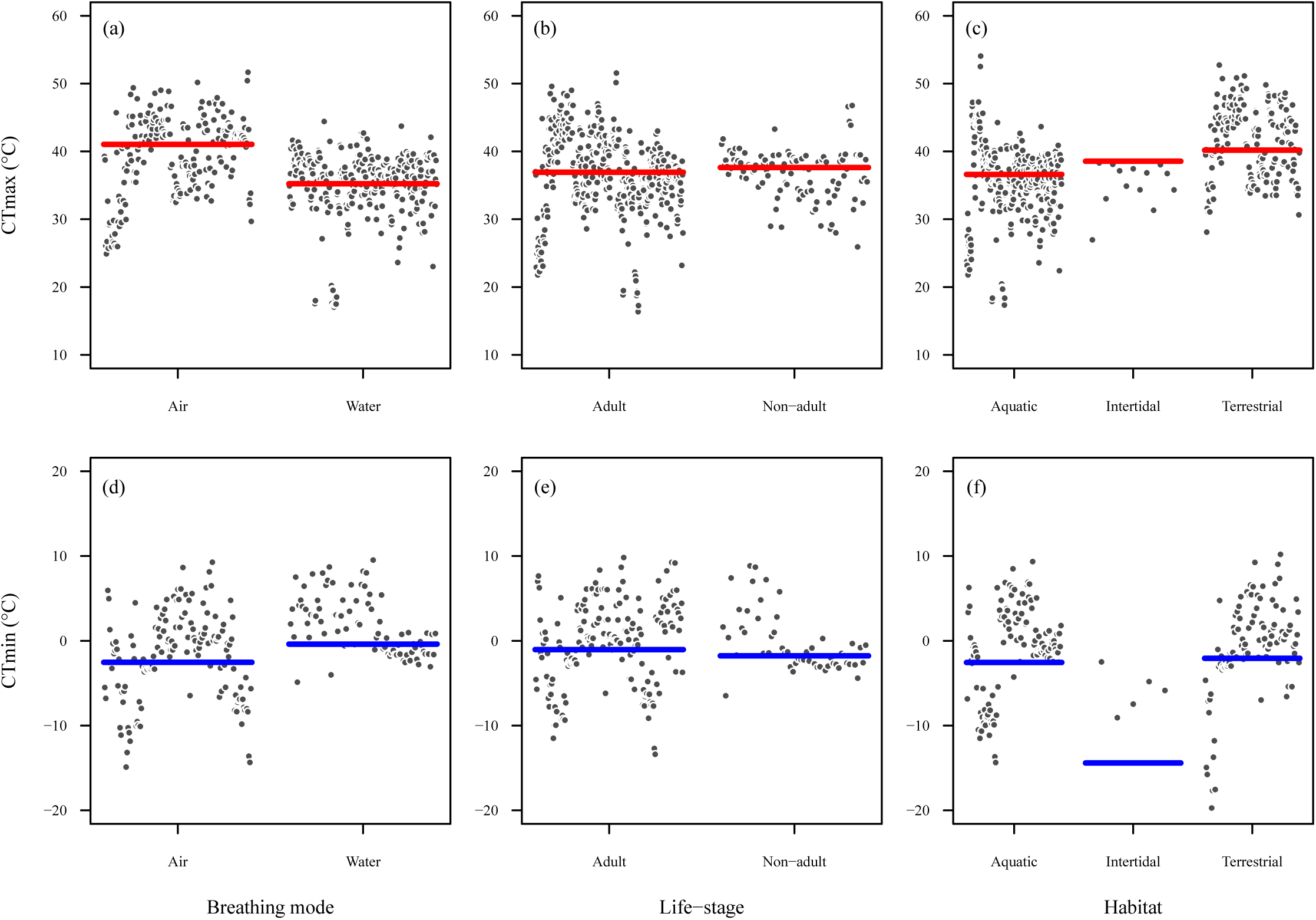
Partial residuals plots showing the predicted effects of breathing mode (a, d), life-stage (b, e) and habitat (c, f) in ectotherms’ CTmax (top, red) and CTmin (bottom, blue). CTmax (a, b, c) and CTmin plots (d, e, f) were based on the model with the highest support shown in Tables 1, 3 that included either breathing mode, life-stage or habitat. Horizontal solid lines are the predicted median of the thermal limits within factor level, conditioned on 2 hours of exposure duration (time) and 45° of absolute latitude. For details on model estimates and significance, see electronic supplementary material Tables S3-S6.

More complex models, testing three interactions (body mass × breathing mode × exposure duration), showed the highest support to explain variations both in CTmax (Table 1 and Table 3) and with some exceptions, in CTmin (Table 2 and Table 4). In general, these models indicate that exposure duration modulates the intensity or even, reverses the direction of the effects of body mass (Figure 3) or genome size (Figure 4). For water-breathers, larger species were found to have a lower CTmax in long-term experimental trials, while the model indicates an opposite effect in short-term trials (Figure 3a,b). For cold tolerance, the three-way interaction with exposure duration was also important for models including body mass and genome size. Here, air-breathers showed improved cold tolerance (lower CTmin values) with increasing genome size, but only for long-term trials (Figure 4d).

**Figure 3.**
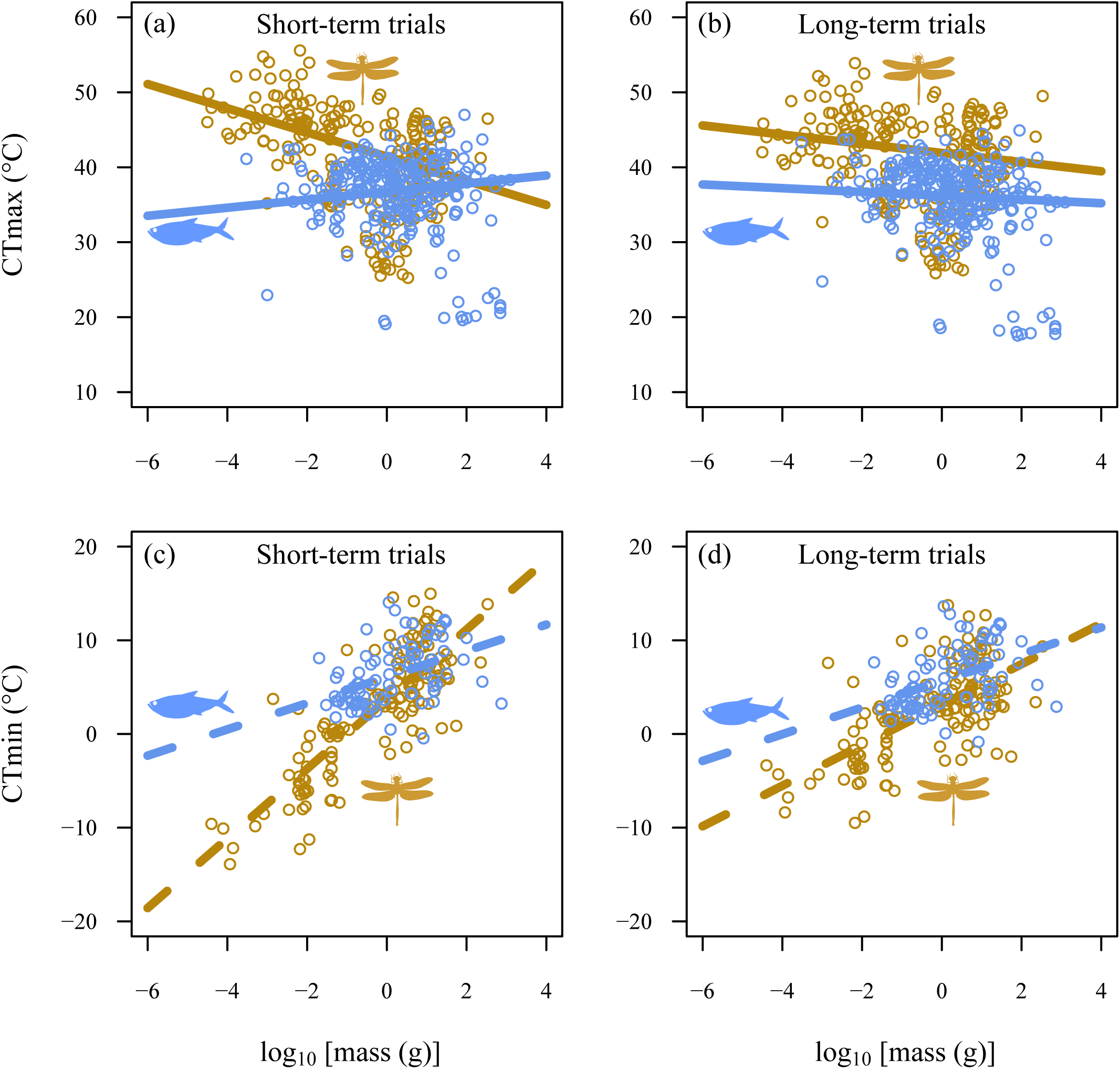
Partial residuals plots showing the interactive effects of body mass and breathing mode (brown and blue for air- and water-breathers, respectively) in ectotherms’ CTmax (top) and CTmin (bottom), for short (1^st^ quartile) and long-term trials (3^rd^ quartile). Predicted lines were based in models with the highest support shown in Tables 1, 2 and based on the median of absolute latitude. For details on model estimates and significance, see electronic supplementary material Tables S3-S4.

**Figure 4.**
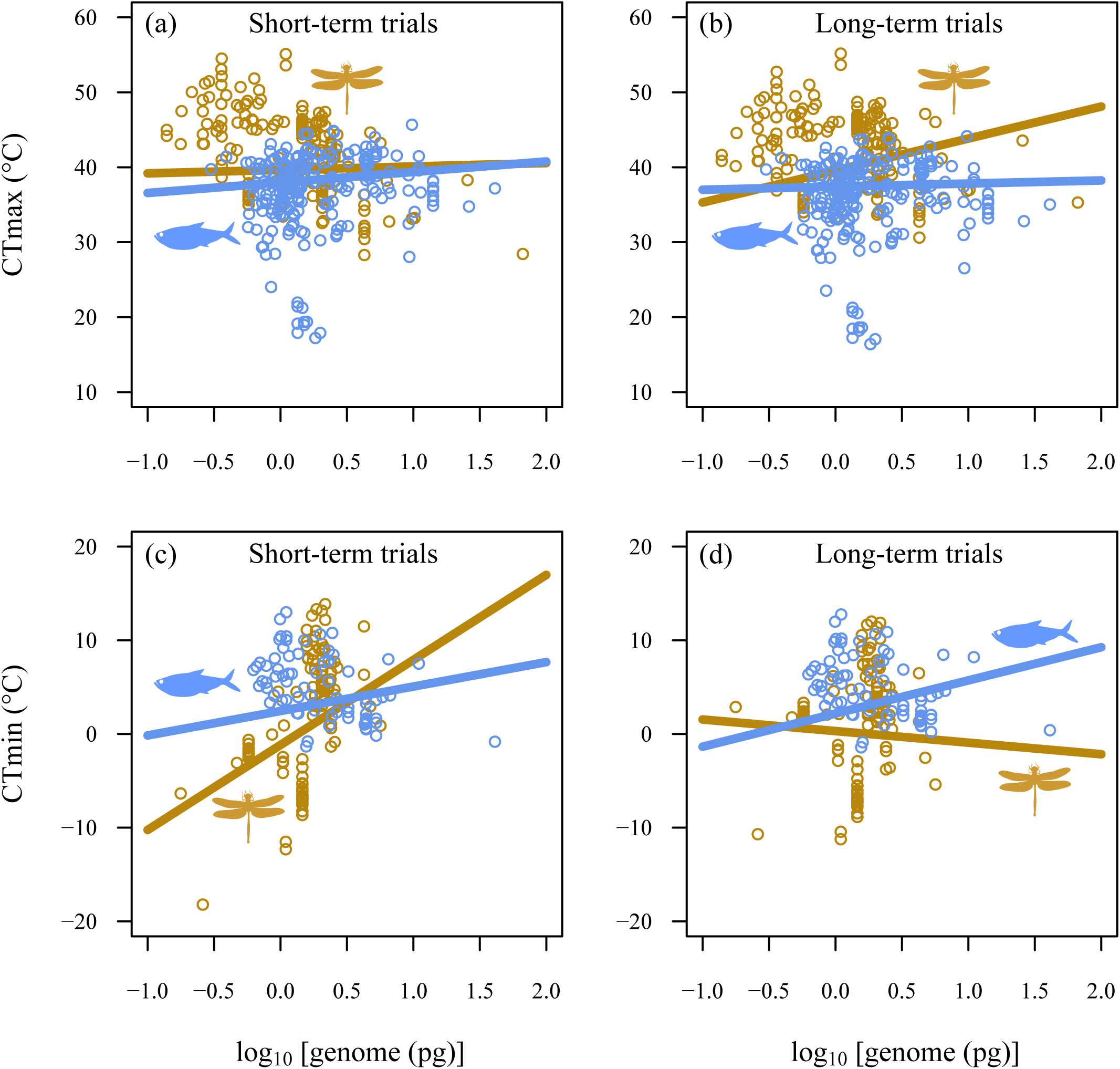
Partial residuals plots showing the interactive effects of genome size and breathing mode (brown and blue for air- and water-breathers, respectively) in ectotherms’ CTmax (top) and CTmin (bottom), for short (1^st^ quartile) and long-term trials (3^rd^ quartile). Predicted lines were based in models with the highest support shown in Tables 3, 4 and based on the median of absolute latitude. For details on model estimates and significance, see electronic supplementary material Tables S5-S6.

Since different numbers of species were included in our analyses on body mass and genome size, the performance of the models cannot be compared directly. We therefore repeated the analyses in a smaller set of species for which information on both body mass and genome size was available. This smaller set included 433 species for CTmax and 190 species for CTmin. These analyses allowed us not only to compare the contributions of body mass and genome size but also test for possible interactions between body mass and genome size. The results of these analyzes were highly consistent with those presented above, that is, models with the highest support, both for the CTmax and CTmin were those that incorporated body mass, genome size, breathing mode and exposure duration. Interestingly, variations in CTmax were mainly driven by those models that considered body mass instead of genome size (Table S7, electronic supplementary material). On the contrary, for the CTmin, the model with the highest support (*w*_i_ = 0.99) was that which considered the three-way interaction of body mass and genome size and exposure duration (Table S9, electronic supplementary material).

## Discussion

Body mass is of fundamental importance for the ecology of ectotherms, governing the rates of energy uptake and energy transformation at the organismal level, with subsequent consequences for species interactions and to the ecosystem functioning. Knowing whether the consequences of global warming are size dependent is therefore central, particularly in light of the ongoing global climatic warming. Here, we have taken a comparative approach to shed light on the relationship between thermal tolerance levels and body mass and genome size in ectotherms. A challenge in such large-scale, comparative studies lies in dealing with the unique evolutionary history of the various species (Spicer & Morley, this issue), as well as dealing with differences in methodology across studies [58,67,68]. Our results show that effects of body mass and genome size on thermal limits (CTmax and CTmin) are context dependent, co-varying to some extent with the evolutionary relationships across species and differing mainly with breathing mode of a species. The methodology was also influential (see also Sunday et al., this issue), as size-dependent differences in thermal limits were magnified in long-term trials.

### Do body mass and genome size relate to thermal limits?

Our results indicate that there is not a simple, straightforward answer as to whether body mass and genome size matters in defining a species’ thermal tolerance or not. If heat tolerance limits arise from insufficient oxygen provisioning to meet demand, and if such oxygen limitation is more likely to occur in larger ectotherms, we would expect heat tolerance to be more impaired in larger-bodied animals. We found such a relationship, but only in our analyses that did not include phylogenetic relationships. Accounting for phylogeny appears to be a more parsimonious explanation for variation in heat tolerance. Still, even when accounting for phylogenetic relationships, we find size dependence of heat tolerance, but this was contingent upon exposure duration and breathing mode, with impaired heat tolerance being more apparent in larger, water-breathers animals during longer trials. Owing to the challenge of underwater gas exchange, water-breathers have been argued to be more susceptible to oxygen-limited heat tolerance [16,21]. The timescale is also important here as stress relates to both its intensity and duration [67]. Heat stress may result in energy deficits, and while energy can be generated either aerobically or anaerobically, anaerobic metabolism is much less efficient and more suitable to deal with acute, short-term energy deficits [69]. For fish, it has been suggested that larger species rely more on anaerobic metabolism when faced with energy deficits [70–72] and if this mass scaling generalizes, this could explain why larger species may be better in coping with short, acute heat stress, but not with prolonged heat stress. Given these considerations, it is perhaps not surprising to find the strongest effects of body mass in prolonged trials on water-breathers.

Heat tolerance was lower in water-breathers compared to air-breathers during prolonged trials when they had larger body sizes, but also when they had larger cell size. The observed effects of genome size can also be interpreted from an oxygen perspective, as smaller genome size is coupled to smaller cell size [73], which can promote a more efficient diffusion of oxygen towards the mitochondria owing to increased membrane surface area to cell volume ratios and shorter diffusion distances [24,44,74,75]. Studies on flies and isopods have shown that warming-induced size reductions are more pronounced under hypoxia [29,74,76], supporting the idea of oxygen shortage setting limits to the size that an animal can attain. This idea also implies that oxygen is unlikely to be limiting in when animals have not yet approached their maximum, species-specific size. As the body mass used here is that of the experimental species, in most cases the specimens used in the experiments will not have represented the upper size classes. This may explain why phylogeny better explains the variation in CTmax: Phylogeny is more likely to covary with the maximum size that a species can attain, but not necessarily with the size of the individuals used in the experiments. Indeed, early, non-adult life stages (i.e. juveniles and larvae) which by definition are not yet fully developed, both show improved heat tolerance with increasing body mass, contrasting with impaired heat tolerance in adults (Figure S9, electronic supplementary material). Along the same line, in a study looking at intraspecific variation in body mass, CTmax improved with body mass in juveniles of a spider species, but deteriorated with size of adults in species of Hemiptera and Collembola [38]. Thus, an oxygen-based mechanism could play a role in heat tolerance but appears to be more relevant for water-breathers and on longer timescales: i.e. exactly those conditions for which a strong warming-induced reduction in body mass has been observed [5].

Unlike heat tolerance, cold tolerance has been suggested to result from depolarization of cell membranes and subsequent cell death [46,77–80], and not from oxygen limitation [81]. Our results also suggests that the mechanisms underpinning CTmin differ from those underpinning of CTmax as the contribution of phylogeny, body mass and genome size to explain variation in CTmin differed when compared to explaining variation in CTmax (Tables S3-S6). Models that consider combined effects of body mass and genome size indicate that this combination better explain variation in CTmin, but not variation in CTmax (see Tables S7-S10, electronic supplementary material). While a small genome size (or small cell size) may enhance oxygen diffusion, it also entails greater costs in keeping membranes polarized [44,82]. Thus, larger cells may be more cost efficient and this could explain why larger genomes can improve cold tolerance. The effect of such an efficiency-based mechanism would likely be more apparent during prolonged trials, and indeed we found that including the interaction between genome size, habitat and exposure duration showed the highest support across all models (Table 4), showing improved cold tolerance in terrestrial animals with larger genome during prolonged trials (Figure 4). In line with these findings, results on the larvae of the pipevine swallowtail *Battus philenor* (Linnaeus 1971) suggested that larger species may have more energy reserves for maintaining metabolism during chill coma, thus explaining their improved cold tolerance [83]. When coupled to lower mass-specific metabolism in larger animals, such an efficiency mechanism would be generally applicable to the whole size range and not only restricted to the largest size classes within a species. This may explain why cold tolerance is most parsimoniously explained by differences in body mass, rather than phylogeny (which is more likely to covary with maximum size but not necessarily with the size of the animals used in the experiment). Interestingly, these patterns for CTmin were more apparent for air-breathers, perhaps because cold tolerance limits in water-breathers are more related to the freezing of water.

### Model fit, phylogenetic correlation structure and covariates

We found evidence of the influence of phylogeny on two fronts. First, both the CTmax and CTmin were phylogenetically structured, displaying high Pagel’s λ (Table S11, electronic supplementary material) and second, those models incorporating phylogeny generally received greater support (especially for CTmax) compared with those where the evolutionary history of the species was considered independent. Also, the Pagel’s λ used in our models (λ = 1) is highly likely to be representative value of the shared evolution of species represented in our database, since all continuous variables, both independent (body mass, genome size, exposure duration and absolute latitude) and dependent (CTmax and CTmin), showed high phylogenetic signals (all λ > 0.7) (Table S11, electronic supplementary material).

The influence of phylogeny on thermal limits is also evident from the contrasting effects of body mass and genome size between models that considered a Brownian or star phylogeny correlation structure (see Figure 1). This indicates that body mass and genome size covary with phylogeny, something which is also evident from the high Pagel’s λ value for body mass and genome size (Table S11, electronic supplementary material). Consequently, incorporating phylogeny already accounts for much of the variation in thermal tolerance, thereby changing the fitted relationship for body mass and genome size. For CTmax, models that included phylogeny always showed greater support, suggesting that the patterns in heat tolerance were more parsimoniously explained by including evolutionary history, possibly because phylogeny better captures the maximum body size which may be causally related to CTmax (see above). For CTmin, models that included the species’ body mass as an independent variable showed greater support when the evolution of the species was assumed as independent (i.e. star phylogeny), possibly because here the actual body size of the experimental individuals is causally related to cold tolerance (see above). Overall, our results confirm earlier findings suggesting that evolutionary history matters for thermal tolerance [84–87], especially for heat tolerance [84,88] and also suggest that this applies to the much larger set of ectotherm species, including insects, crustaceans, fish, amphibians and reptiles. Our results also point out the importance of including, mainly methodological aspects as covariates into modelling (see also Sunday et al., this issue). Of the four methodological aspects evaluated in the preliminary models (absolute latitude, exposure duration, acclimation time and origin), latitude and exposure duration were consistently included in the best-fitted models. The absolute latitude of the site where animals were collected consistently shifted thermal windows, impairing the heat tolerance and improving the cold tolerance at higher latitudes (Figure S7). While their effects were not the focus of our analyses, these results reinforce the clear patterns of thermal tolerance across latitudinal gradients documented in the literature [12] (see also Sunday et al. this issue). The exposure duration was also consistently included in the best-fitted models, with long-term trials consistently reducing CTmax. This indicates that methodological variation explains a significant part of the variation in CTmax and adding exposure duration as covariate may help to reveal more clearly the effects of other factors, such as that of latitude [67]. Furthermore, the inclusion of exposure duration as a covariate has direct biological meaning as tolerance to high-temperature conditions is time-dependent [67].

## Conclusion

In conclusion, for CTmax we found that support for the oxygen limitation hypothesis was limited to long-term trials in larger-bodied water-breathers. For CTmin, we found improved cold tolerance for air-breather animals with larger genomes, again when considering long-term trials. Coping with thermal stress on long timescales requires sustained energy generation. Long-term heat resistance appears to be enhanced in smaller bodied, water-breathing species, possibly as this enables a higher capacity to generate energy aerobically and efficiently. On the other hand, long-term cold resistance appears to be enhanced in species with a larger body mass and cell size (i.e. more energy reserves and lower energetic costs), which appeared especially important for air-breathers. Incorporating the exposure duration of the experimental trials can reveal body and genome size-dependence of thermal tolerance, with body size being more important for CTmax and water-breathers and genome size being more important for CTmin and air-breathers. Our results highlight the importance of accounting for phylogeny and exposure duration. Especially when considering long-term trials these effects are more in line with the warming-induced reduction in body mass observed during long-term rearing experiments [5] and over past extinctions [8]. Explicitly incorporating time scale may thus hold the key to resolve discrepancies between short-term trials that do not always find evidence for oxygen limitation and the results of long-term laboratory and field studies that do suggest a role for oxygen limitation.

## Supporting information

Electronic supplementary material

## Ethics

This study did not require ethical approval because no animal was handled and no fieldwork was carried out. Analyzed data were obtained from publicly published studies.

## Data accessibility

The dataset supporting this article have been uploaded as part of the electronic supplementary material.

## Authors’ contributions

F.P.L. extracted the data from the articles; F.P.L. conducted the statistical analyses and prepared figures with inputs from W.C.E.P.V. and P.C; F.P.L. and W.C.E.P.V. wrote the first version of the manuscript with inputs from P.C; All authors contributed and approved the final version of the MS.

## Acknowledgements

We thank D. Sánchez–Fernández, L. Viegas, L.M. Gutierrez–Pesquera and A. Barría for kindly provide body mass data of the experimental animals. We also thank Jennifer Sunday and two anonymous reviewers for helpful comments and suggestions that greatly improved this manuscript.

