## Supplementary material for "Scaling of thermal tolerance with body mass and genome size in ectotherms: A comparison between water-and air-breathers": Electronic supplementary material

Supplementary figures: Figures S1-S9
Supplementary tables: Tables S1-S11

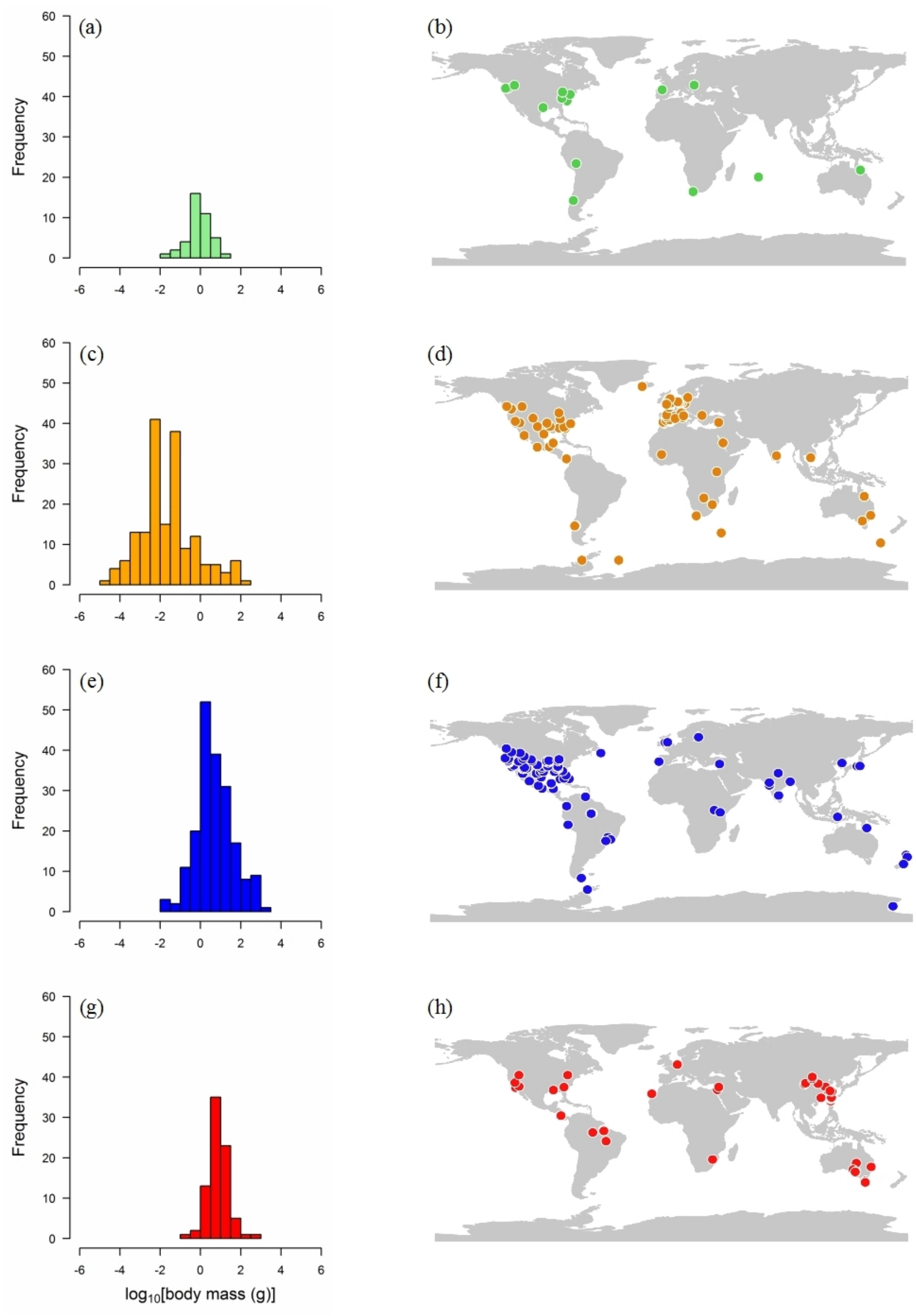


**Figure S1.** Frequency per body mass (left hand) and geographic distribution of species (right hand) used in thermal tolerance experiments in amphibians (a, b), arthropods (c, d), fish (e, f) and reptiles (g, h).

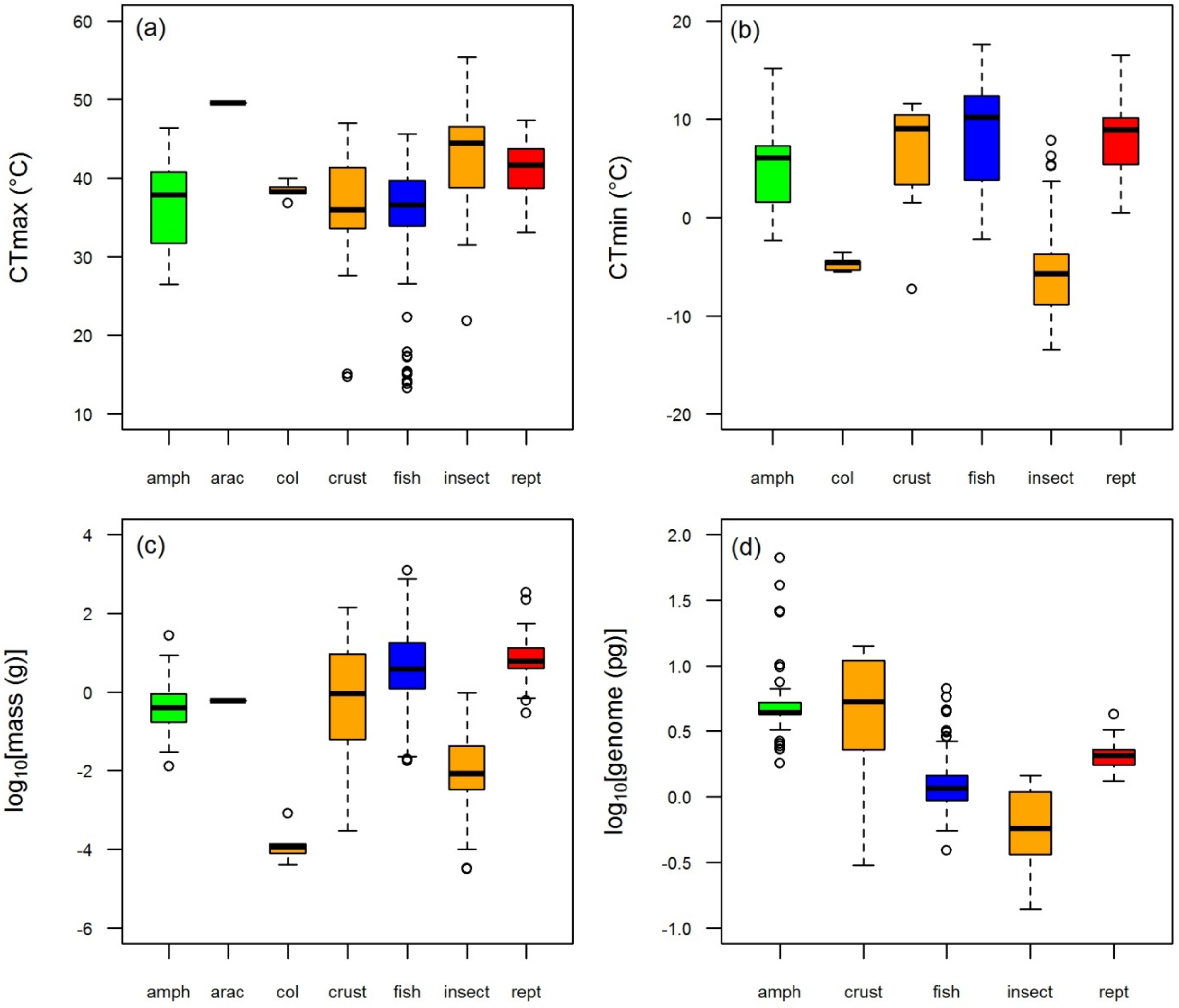


**Figure S2**. CTmax (a), CTmin (b), body mass (c) and genome size (d) in amphibians (amph), arachnids (arac), collembola (col), crustaceans (crust) fish, insect and reptiles (rept) species used in our analyses.

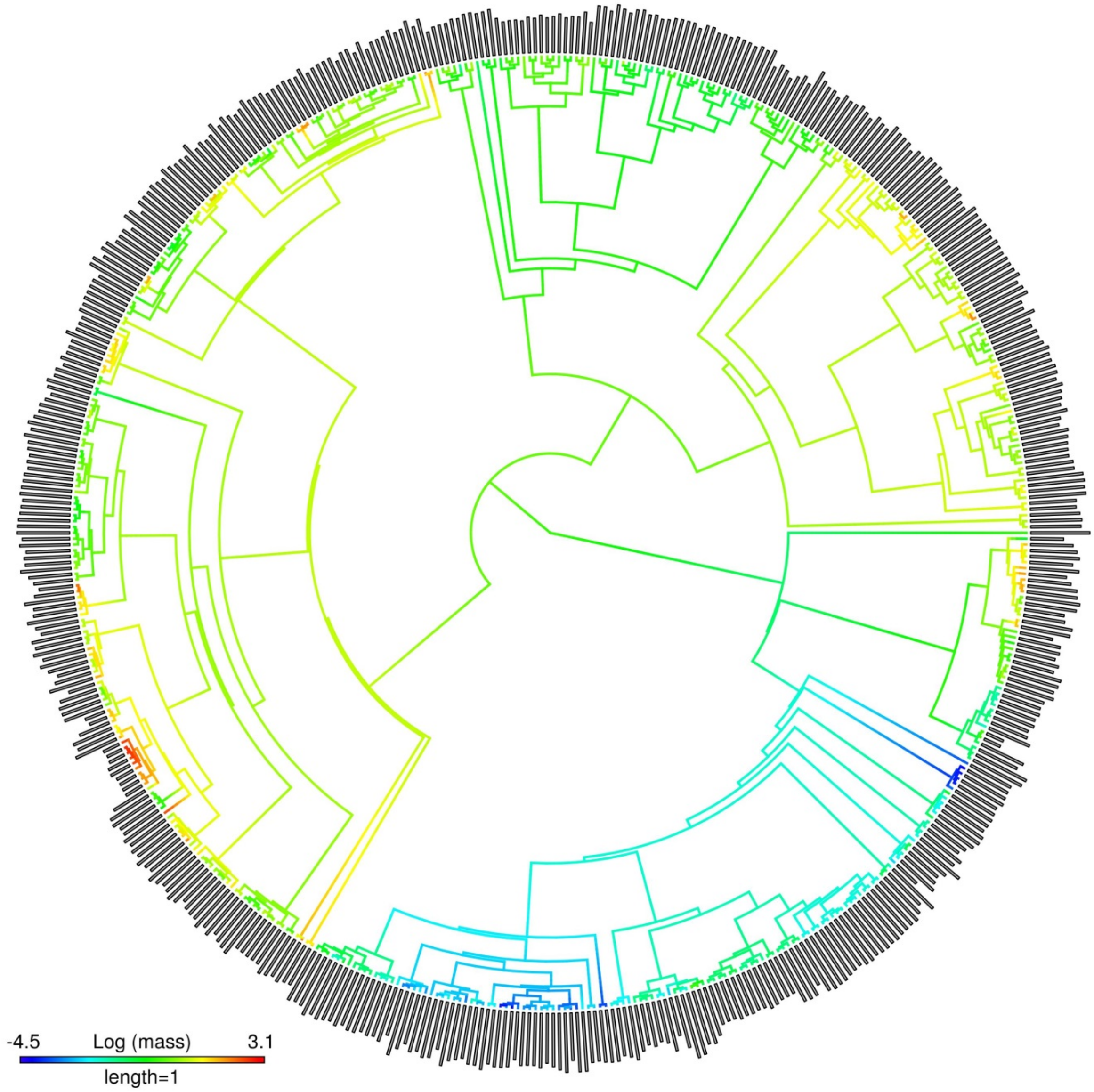


27

28 **Figure S3**. Phylogenetic tree displaying the relation between CTmax (outer grey bars) and body mass (gradient in
29 the branch' colors) of ectotherms' species used in this study (N = 510).

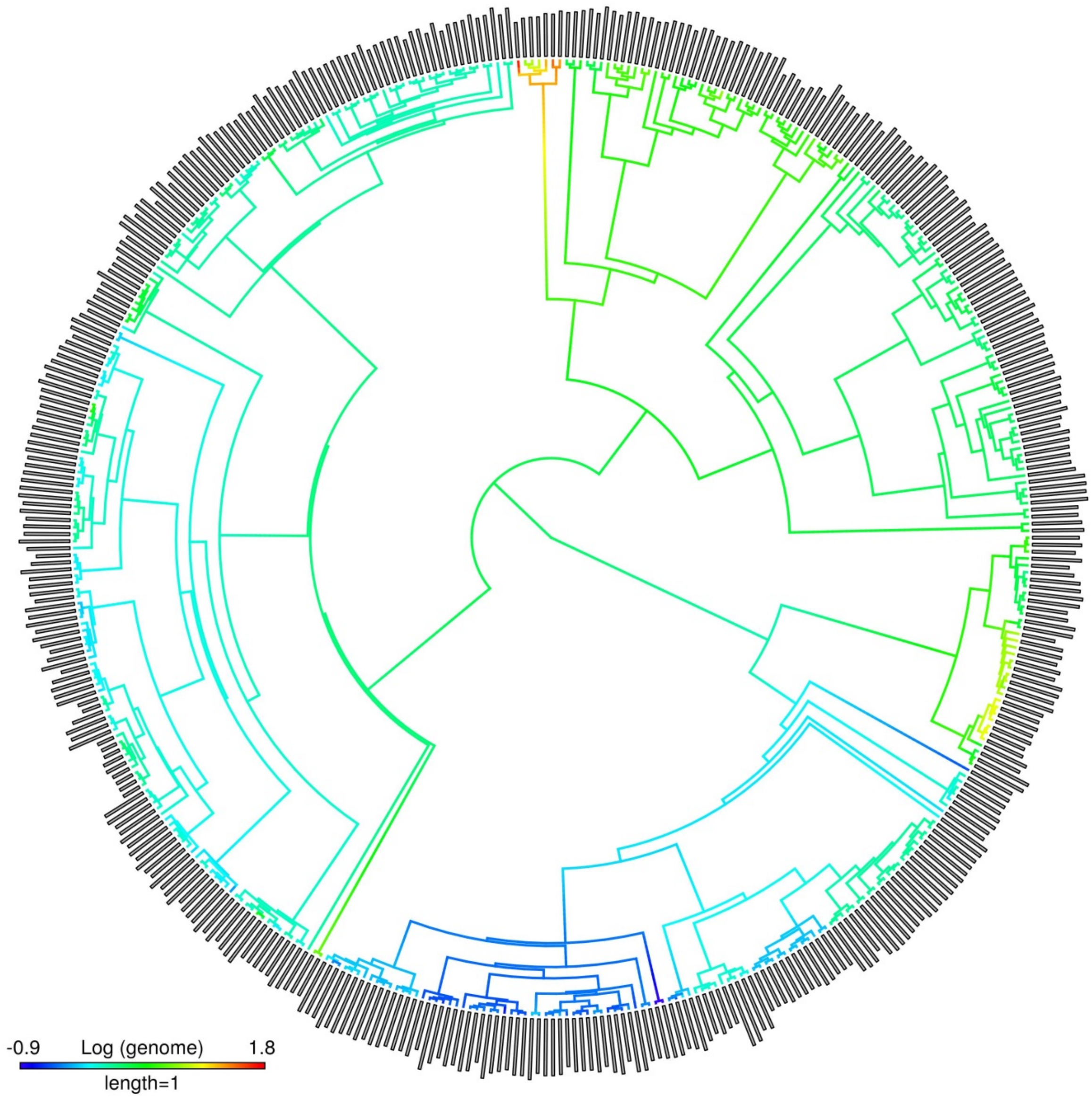


**Figure S4.** Phylogenetic tree displaying the relation between CTmax (outer grey bars) and genome size (gradient in the branch' colors) of ectotherms' species used in this study (N = 433).

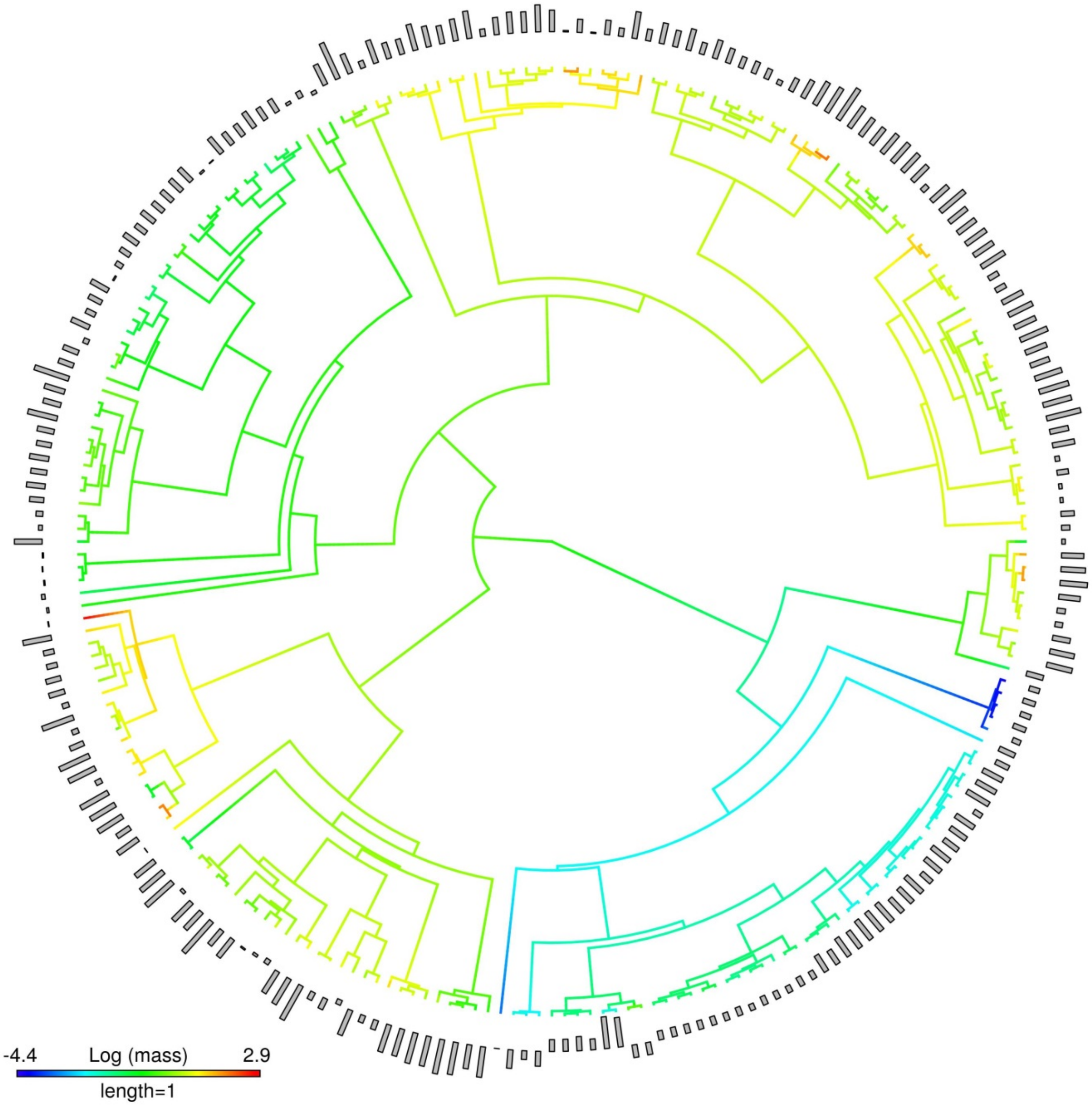


**Figure S5.** Phylogenetic tree displaying the relation between CTmin (outer grey bars) and genome size (gradient in the branch' colors) of ectotherms' species (N = 232).

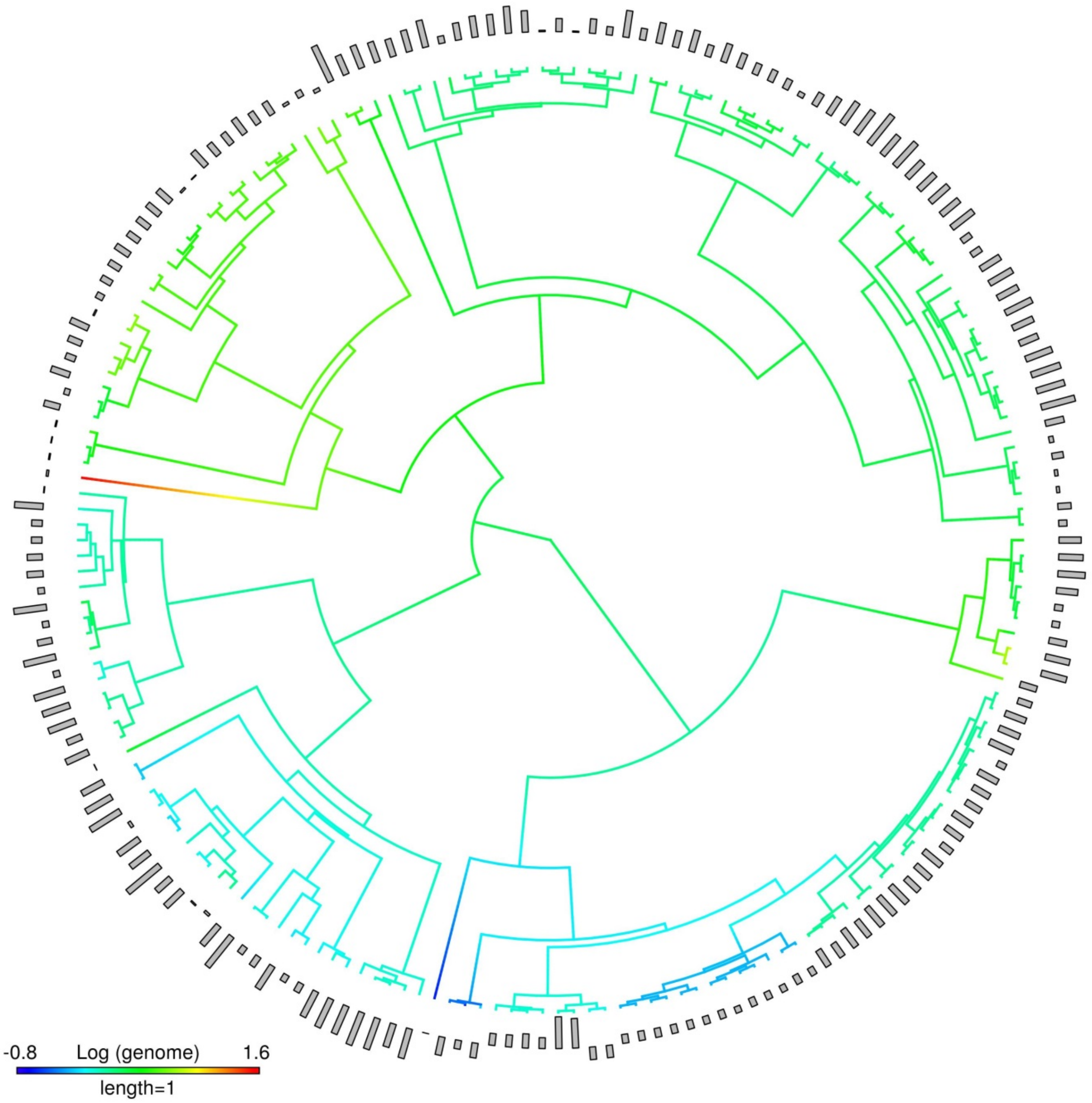


**Figure S6.** Phylogenetic tree displaying the relation between CTmin (outer grey bars) and genome size (gradient in the branch' colors) of ectotherms' species used in this study (N = 190).

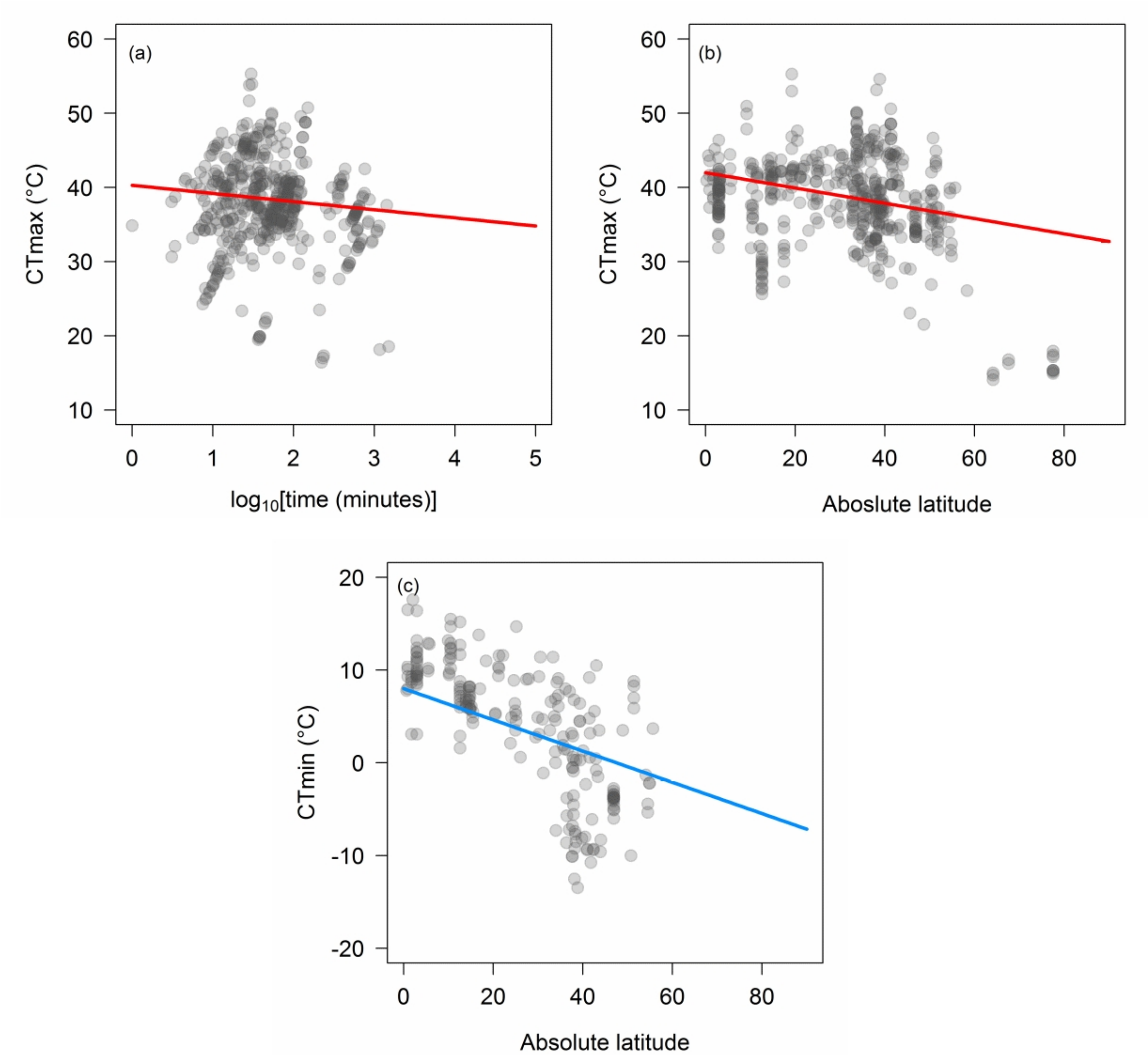


**Figure S7**. Partial residuals plots showing the effects (solid lines) of $\log_{10}$–transformed time (a) and absolute latitude in ectotherms' CTmax (top, red line) and CTmin (bottom, blue line). Predicted lines for CTmax were based in the model with the highest support; for CTmax model 1 and for CTmin model 19 (see Table S1). For details on model' estimates and significance, see Table S2.

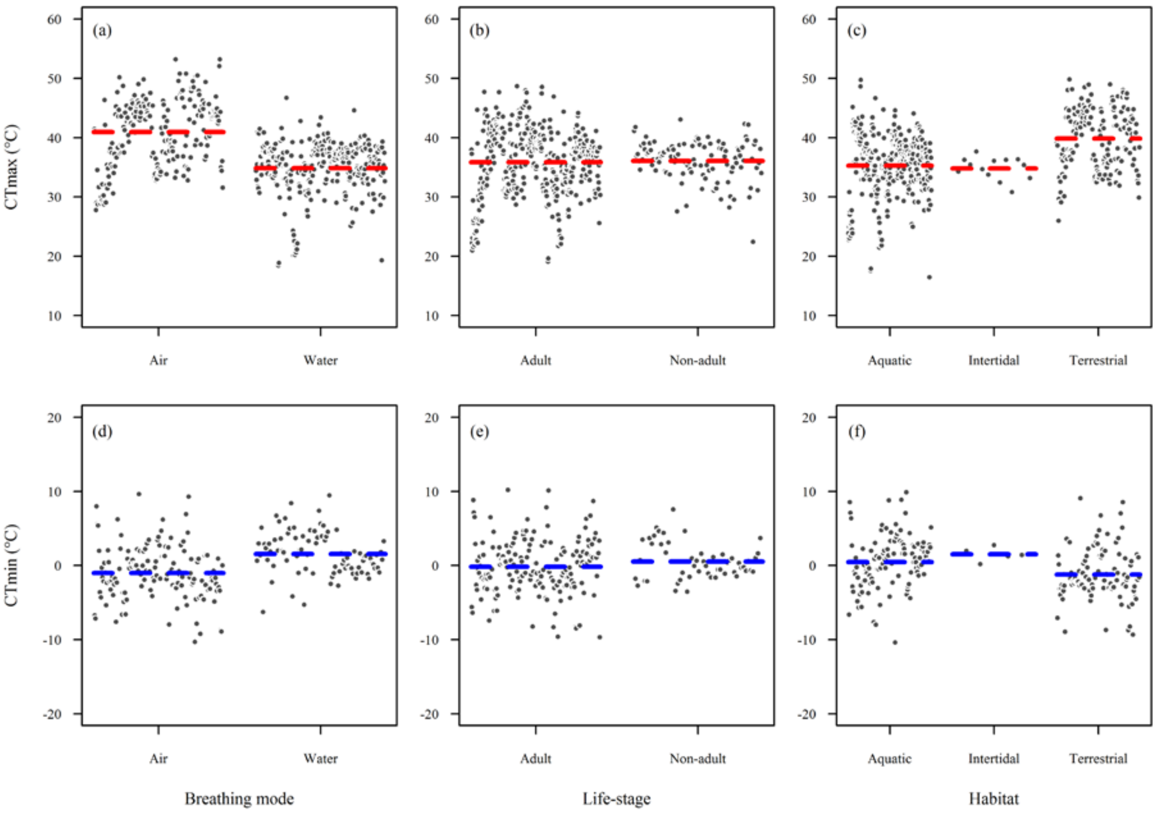


**Figure S8.**Partial residuals plots showing the predicted effects of breathing mode (a, d), life-stage (b, e) and habitat (c, f) in ectotherms' CTmax (top, red) and CTmin (bottom, blue). CTmax (a, b, c) and CTmin plots (d, e, f) were based in that included all covariates, but not phylogeny ($\lambda = 0$). Horizontal dashed lines are the predicted median of the thermal limits within factor level, conditioned on 2 hours of exposure duration (time) and 45° of absolute latitude. For details on model estimates and significance, see electronic supplementary material Tables S3-S6.

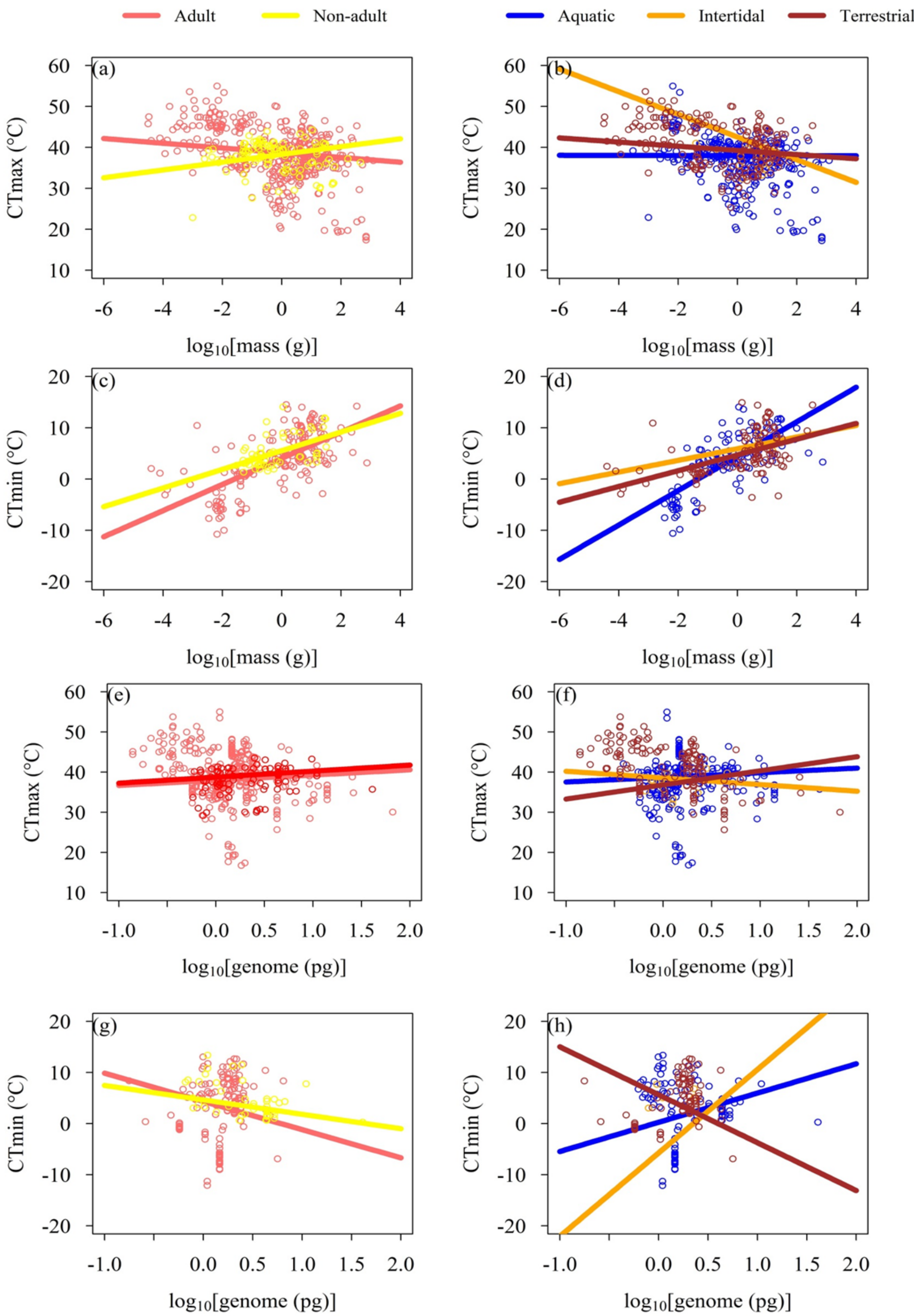


**Figure S9**. Partial residuals plots showing the interactive effects (solid lines) of body mass with life-stage (a, c), habitat (b, d), and genome size life-stage (e, g), habitat (f, h) in ectotherms' CTmax and CTmin. Predicted lines were based on models Table 1 and Table 2. Predicted lines were based at median values of two covariates (absolute latitude and/or time).

**Table S1**. Selection results of PGLSs models to explain variation in ectotherms' CTmax and CTmin as function of exposure duration (Time), absolute latitude (Lat) acclimation time ($t_a$*:* $\log_{10}$–transformed [acclimation time + 1]) and origin (laboratory or field). Pagel's lambda ($\lambda$) denotes correlation structure used ($\lambda$ = 0, star phylogeny and $\lambda$ = 1, Brownian phylogeny). Number of parameters (*k*), corrected Akaike's information criterion (AICc), difference in AICc respect to the model with highest support (ΔAICc), log-likelihood (*LL*) and the Akaike's weights ($w_i$) are mentioned for each model. (N) indicates the number of species used in each model.

| **Models** | **λ** | ***k*** | **AICc** | **ΔAICc** | ***LL*** | $w_i$ |
|---|---|---|---|---|---|---|
| CTmax (N = 478) | | | | | | |
| **1. Lat + Time** | **1** | **4** | **2781.19** | **0** | **-1386.55** | **0.71** |
| **2. Lat × Time** | **1** | **5** | **2782.99** | **1.80** | **-1386.43** | **0.29** |
| 3. Lat | 1 | 3 | 2797.65 | 16.47 | -1395.8 | 0.00 |
| 4. $t_a$ × Origin + Lat | 1 | 6 | 2799.73 | 18.54 | -1393.77 | 0.00 |
| 5. $t_a$ × Origin + Time | 1 | 6 | 2832.17 | 50.98 | -1409.99 | 0.00 |
| 6. Time | 1 | 3 | 2834.48 | 53.29 | -1414.21 | 0.00 |
| 7. $t_a$ | 1 | 3 | 2844.15 | 62.97 | -1419.05 | 0.00 |
| 8. $t_a$ × Origin | 1 | 5 | 2846.15 | 64.97 | -1418.01 | 0.00 |
| 9. Origin | 1 | 3 | 2851.3 | 70.11 | -1422.62 | 0.00 |
| 10. Lat × Time | 0 | 5 | 3075.48 | 294.29 | -1532.67 | 0.00 |
| 11. Lat + Time | 0 | 4 | 3082.69 | 301.50 | -1537.3 | 0.00 |
| 12. Lat | 0 | 3 | 3084.63 | 303.44 | -1539.29 | 0.00 |
| 13. $t_a$ × Origin + Lat | 0 | 6 | 3089.72 | 308.53 | -1538.77 | 0.00 |
| 14. $t_a$ * Origin + Time | 0 | 6 | 3132.91 | 351.73 | -1560.37 | 0.00 |
| 15. Time | 0 | 3 | 3134.57 | 353.39 | -1564.26 | 0.00 |
| 16. $t_a$ | 0 | 3 | 3134.73 | 353.54 | -1564.34 | 0.00 |
| 17. $t_a$ × Origin | 0 | 5 | 3138.58 | 357.40 | -1564.23 | 0.00 |
| 18. Origin | 0 | 3 | 3140.81 | 359.63 | -1567.38 | 0.00 |
| CTmin (N = 208) | | | | | | |
| **19. Lat** | **1** | **3** | **1149.12** | 0.00 | **-571.50** | **0.51** |
| **20. Lat × Time** | **1** | **5** | **1150.17** | 1.05 | **-569.94** | **0.30** |
| **21. Lat + Time** | **1** | **4** | **1151.12** | 2.00 | **-571.46** | **0.19** |
| 22. $t_a$ | 1 | 3 | 1217.30 | 68.18 | -605.59 | 0.00 |
| 23. Time | 1 | 3 | 1237.63 | 88.51 | -615.75 | 0.00 |
| 24. Lat | 0 | 3 | 1253.36 | 104.24 | -623.62 | 0.00 |
| 25. Lat + Time | 0 | 4 | 1253.73 | 104.61 | -622.77 | 0.00 |
| 26. Lat × Time | 0 | 5 | 1255.66 | 106.54 | -622.68 | 0.00 |
| 27. $t_a$ | 0 | 3 | 1399.32 | 250.20 | -696.60 | 0.00 |
| 28. Time | 0 | 3 | 1432.71 | 253.59 | -698.30 | 0.00 |

**Table S2.** Summary outputs of preliminary phylogenetic generalized least squares (PGLSs) models with the highest support ($w_i$) for CTmax and CTmin analyses enlisted in Table S1. Pagel's lambda (λ) denotes correlation structure used (λ = 0, star phylogeny and λ = 1, Brownian phylogeny). (N) indicates the number of species used in each model.

| **Model** | **Variable** | **Estimate** | **SE** | ***P*-value** |
|---|---|---|---|---|
| CTmax (N = 478) | | | | |
| Lat + Time (λ = 1)<br>Table S1; Model 1 | Intercept | 43.791 | 13.863 | 0.0017 |
| | Lat | -0.103 | 0.013 | < **0.0001** |
| | Time | -1.097 | 0.253 | < **0.0001** |
| Lat × Time (λ = 1)<br>Table S1; Model 2 | Intercept | 43.404 | 13.896 | 0.0019 |
| | **Lat** | **-0.088** | **0.032** | **0.0069** |
| | Time | -0.904 | 0.470 | 0.0551 |
| | Lat × Time | -0.007 | 0.015 | 0.6259 |
| CTmin (N = 208) | | | | |
| Lat (λ = 1)<br>Table S1; Model 19 | Intercept | 8.00 | 9.50 | 0.4004 |
| | **Lat** | **-0.17** | **0.02** | < **0.0001** |
| Lat × Time (λ = 1)<br>Table S1; Model 20 | Intercept | 6.80 | 9.56 | 0.4780 |
| | Lat | -0.07 | 0.06 | 0.2342 |
| | Time | 0.57 | 0.61 | 0.3527 |
| | Lat × Time | -0.05 | 0.03 | 0.0840 |
| Lat + Time (λ = 1)<br>Table S1; Model 21 | Intercept | 8.26 | 9.57 | 0.3888 |
| | **Lat** | **-0.17** | **0.02** | < **0.0001** |
| | Time | -0.13 | 0.46 | 0.7832 |

**Table S3.** Summary outputs for the phylogenetic generalized least squares (PGLSs) model with the highest support ($w_i$) for CTmax analysis enlisted in Table 1, main text. Model considered as covariate the absolute latitude (Lat) of animal collection. Pagel's lambda λ = 1 denotes correlation structure used (Brownian phylogeny).

| **Model** | **Variable** | **Estimate** | **SE** | ***P*-value** |
|---|---|---|---|---|
| Body mass × Breathing mode × Time + Lat (λ = 1) | Intercept | 43.48 | 13.93 | 0.0019 |
| | **Body mass** | **-3.74** | **0.85** | **0.0000** |
| | Water | -2.46 | 2.08 | 0.2374 |
| | Time | 0.79 | 1.14 | 0.4877 |
| | **Lat** | **-0.09** | **0.01** | **0.0000** |
| | **Body mass × Water** | **5.96** | **1.04** | **0.0000** |
| | **Body mass × Time** | **1.62** | **0.47** | **0.0007** |
| | Water × Time | -1.68 | 1.18 | 0.1536 |
| | **Body mass × Water × Time** | **-2.90** | **0.57** | **0.0000** |

**Table S4.** Summary outputs for the phylogenetic generalized least squares (PGLSs) model with the highest support ($w_i$) for CTmin analysis enlisted in Table 2, main text. Model considered as covariate the absolute latitude (Lat) of animal collection. Pagel's lambda λ = 0 denotes correlation structure used (star phylogeny).

| **Model** | **Variable** | **Estimate** | **SE** | ***P*-value** |
|---|---|---|---|---|
| Body mass × Breathing mode × Time + Lat (λ = 0) | Intercept | 11.86 | 1.33 | 0.0000 |
| | **Body mass** | **7.75** | **0.77** | **0.0000** |
| | Water | 1.92 | 1.82 | 0.2933 |
| | Time | -1.14 | 0.80 | 0.1554 |
| | **Lat** | **-0.24** | **0.02** | **0.0000** |
| | **Body mass × Water** | **-6.42** | **1.75** | **0.0003** |
| | **Body mass × Time** | **-3.04** | **0.44** | **0.0000** |
| | Water × Time | 0.34 | 1.09 | 0.7560 |
| | **Body mass × Water × Time** | **3.09** | **0.98** | **0.0019** |

**Table S5.** Summary outputs for phylogenetic generalized least squares (PGLSs) models with the highest support ($w_i$) for CTmax analysis enlisted in Table 3, main text. Models considered as covariates the exposure duration (Time) and/or absolute latitude (Lat) of animal collection. Pagel's lambda $\lambda = 1$ denotes correlation structure used (Brownian phylogeny).

| **Model** | **Variable** | **Estimate** | **SE** | ***P*-value** |
|---|---|---|---|---|
| Genome size × Breathing mode × Time + Lat ($\lambda$ = 1); AICc = 2502.36; $w_i$ = 0.36; ΔAICc = 0.00 | Intercept | 42.88 | 13.78 | 0.0020 |
| | **Genome size** | **-8.32** | **4.01** | **0.0386** |
| | Water | -0.58 | 2.18 | 0.7899 |
| | Time | -0.14 | 0.95 | 0.8847 |
| | **Lat** | **0.09** | **0.01** | **0.0000** |
| | **Genome size × Water** | **12.00** | **4.57** | **0.0089** |
| | **Genome size × Time** | **6.54** | **2.23** | **0.0035** |
| | Water × Time | -0.82 | 0.99 | 0.4126 |
| | **Genome size × Water × Time** | **-8.24** | **2.59** | **0.0016** |
| Life-stage + Time + Lat ($\lambda$ = 1); AICc = 2503.91; $w_i$ = 0.16; ΔAICc = 1.55 | Intercept | 43.40 | 13.78 | 0.0017 |
| | Non-adult | 0.83 | 0.43 | 0.0547 |
| | **Lat** | **-0.10** | **0.01** | **0.0000** |
| | **Time** | **-0.95** | **0.27** | **0.0005** |
| Breathing mode + Time + Lat ($\lambda$ = 1); AICc = 2504.30; $w_i$ = 0.13; ΔAICc = 1.95 | Intercept | 45.13 | 13.79 | 0.0012 |
| | Water | -2.08 | 1.15 | 0.0692 |
| | **Lat** | **-0.09** | **0.01** | **0.0000** |
| | **Time** | **-1.08** | **0.26** | **0.0000** |

**Table S6.** Summary outputs for the phylogenetic generalized least squares (PGLSs) model with the highest support ($w_i$) for CTmin analysis enlisted in Table 4, main text. Model considered as covariate the absolute latitude (Lat) of animal collection. Pagel's lambda λ = 1 denotes correlation structure used (Brownian phylogeny).

| **Model** | **Variable** | **Estimate** | **SE** | ***P*-value** |
|---|---|---|---|---|
| Genome size × Habitat × Time + Lat (λ = 1) | Intercept | 7.62 | 9.23 | 0.4100 |
| | Genome size | 3.09 | 5.11 | 0.5471 |
| | Intertidal | -8.16 | 8.19 | 0.3204 |
| | Terrestrial | 0.40 | 3.89 | 0.9185 |
| | Time | -0.17 | 0.59 | 0.7752 |
| | **Lat** | **-0.24** | **0.02** | **0.0000** |
| | Genome size × Intertidal | 59.07 | 32.33 | 0.0694 |
| | Genome size × Terrestrial | 18.21 | 10.88 | 0.0961 |
| | Genome size × Time | 1.95 | 2.74 | 0.4777 |
| | Intertidal × Time | 0.62 | 4.59 | 0.8930 |
| | Terrestrial × Time | 1.70 | 1.71 | 0.3217 |
| | Genome size × Intertidal × Time | -34.54 | 23.52 | 0.1438 |
| | **Genome size × Terrestrial × Time** | **-16.45** | **5.07** | **0.0014** |

**Table S7.** Selection results for phylogenetic generalized least squares (PGLSs) models to explain variation in ectotherms' CTmax (N = 433 species) as function of $log_{10}$–transformed body mass, $log_{10}$–transformed genome size, breathing mode (air and water), life-stage (adults and non-adults), habitat (aquatic, intertidal and terrestrial) and all their possible interactions. Each model was assessed using as covariates the exposure duration (Time) and/or absolute latitude (Lat) of animal collection. Pagel's lambda (λ) denotes correlation structure used (λ = 0, star phylogeny and λ = 1, Brownian phylogeny). Number of parameters (*k*), corrected Akaike's information criterion (AICc), difference in AICc respect to the model with highest support (ΔAICc) and the Akaike's weights ($w_i$) are mentioned for each model.

| **Models** | | ***k*** | **AICc** | **ΔAICc** | $w_i$ |
|---|---|---|---|---|---|
| 0. Covariates | λ = 1 + Lat + Time | 4 | 2505.59 | 17.18 | 0.00 |
| | λ = 0 + Lat + Time | 4 | 2759.27 | 270.86 | 0.00 |
| | λ = 1 + Lat | 3 | 2521.96 | 33.54 | 0.00 |
| | λ = 1 + Time | 3 | 2555.30 | 66.88 | 0.00 |
| 1. Body mass | λ = 1 + Lat + Time | 5 | 2506.74 | 18.32 | 0.00 |
| | λ = 0 + Lat + Time | 5 | 2630.85 | 142.43 | 0.00 |
| | λ = 1 + Lat | 4 | 2523.46 | 35.05 | 0.00 |
| | λ = 1 + Time | 4 | 2554.45 | 66.03 | 0.00 |
| 2. Breathing Mode | λ = 1 + Lat + Time | 5 | 2504.30 | 15.89 | 0.00 |
| | λ = 0 + Lat + Time | 5 | 2644.82 | 156.40 | 0.00 |
| | λ = 1 + Lat | 4 | 2519.78 | 31.37 | 0.00 |
| | λ = 1 + Time | 4 | 2551.15 | 62.74 | 0.00 |
| 3. Life-stage | λ = 1 + Lat + Time | 5 | 2503.91 | 15.50 | 0.00 |
| | λ = 0 + Lat + Time | 5 | 2758.50 | 270.08 | 0.00 |
| | λ = 1 + Lat | 4 | 2513.96 | 25.54 | 0.00 |
| | λ = 1 + Time | 4 | 2551.56 | 63.15 | 0.00 |
| 4. Habitat | λ = 1 + Lat + Time | 6 | 2509.48 | 21.07 | 0.00 |
| | λ = 0 + Lat + Time | 6 | 2718.37 | 229.95 | 0.00 |
| | λ = 1 + Lat | 5 | 2525.79 | 37.38 | 0.00 |
| | λ = 1 + Time | 5 | 2558.64 | 70.22 | 0.00 |
| 5. Body mass × Breathing Mode | λ = 1 + Lat + Time | 7 | 2506.49 | 18.08 | 0.00 |
| | λ = 0 + Lat + Time | 7 | 2576.24 | 87.82 | 0.00 |
| | λ = 1 + Lat | 6 | 2522.65 | 34.23 | 0.00 |
| | λ = 1 + Time | 6 | 2550.92 | 62.50 | 0.00 |
| 6. Body mass × Life-stage | λ = 1 + Lat + Time | 7 | 2501.03 | 12.62 | 0.00 |
| | λ = 0 + Lat + Time | 7 | 2624.92 | 136.51 | 0.00 |
| | λ = 1 + Lat | 6 | 2513.35 | 24.93 | 0.00 |
| | λ = 1 + Time | 6 | 2547.48 | 59.06 | 0.00 |
| 7. Body mass × Habitat | λ = 1 + Lat + Time | 9 | 2514.86 | 26.45 | 0.00 |
| | λ = 0 + Lat + Time | 9 | 2609.24 | 120.83 | 0.00 |
| | λ = 1 + Lat | 8 | 2531.23 | 42.82 | 0.00 |
| | λ = 1 + Time | 8 | 2558.98 | 70.57 | 0.00 |
| **8. Body mass × Breathing mode × Time** | **λ = 1 + Lat** | **10** | **2488.41** | **0.00** | **0.57** |
| | λ = 0 + Lat | 10 | 2574.01 | 85.60 | 0.00 |
| **9. Body mass × Life-stage × Time** | **λ = 1 + Lat + Time** | **10** | **2490.42** | **2.00** | **0.21** |
| | λ = 0 + Lat + Time | 10 | 2621.53 | 133.12 | 0.00 |
| 10. Body mass × Habitat × Time | λ = 1 + Lat + Time | 14 | 2499.39 | 10.98 | 0.00 |
| | λ = 0 + Lat + Time | 14 | 2608.21 | 119.79 | 0.00 |

**Table S7 continued**

| **Models** | | ***k*** | **AICc** | **ΔAICc** | $\boldsymbol{w_i}$ |
|---|---|---|---|---|---|
| 11. Genome size | λ = 1 + Lat + Time | 5 | 2506.18 | 17.76 | 0.00 |
| | λ = 0 + Lat + Time | 5 | 2731.05 | 242.64 | 0.00 |
| | λ = 1 + Lat | 4 | 2521.94 | 33.52 | 0.00 |
| | λ = 1 + Time | 4 | 2554.25 | 65.84 | 0.00 |
| 12. Genome size × Breathing mode | λ = 1 + Lat + Time | 7 | 2506.43 | 18.01 | 0.00 |
| | λ = 0 + Lat + Time | 7 | 2599.70 | 111.28 | 0.00 |
| | λ = 1 + Lat | 6 | 2521.26 | 32.85 | 0.00 |
| | λ = 1 + Time | 6 | 2551.27 | 62.85 | 0.00 |
| 13. Genome size × Life-stage | λ = 1 + Lat + Time | 7 | 2506.65 | 18.24 | 0.00 |
| | λ = 0 + Lat + Time | 7 | 2715.88 | 227.47 | 0.00 |
| | λ = 1 + Lat | 6 | 2515.68 | 27.27 | 0.00 |
| | λ = 1 + Time | 6 | 2552.49 | 64.08 | 0.00 |
| 14. Genome size × Habitat | λ = 1 + Lat + Time | 9 | 2511.67 | 23.25 | 0.00 |
| | λ = 0 + Lat + Time | 9 | 2669.46 | 181.04 | 0.00 |
| | λ = 1 + Lat | 8 | 2527.59 | 39.17 | 0.00 |
| | λ = 1 + Time | 8 | 2553.99 | 65.58 | 0.00 |
| 15. Genome size × Breathing mode × Time | λ = 1 + Lat | 10 | 2502.36 | 13.94 | 0.00 |
| | λ = 0 + Lat | 10 | 2590.26 | 101.84 | 0.00 |
| 16. Genome size × Life-stage × Time | λ = 1 + Lat | 10 | 2510.08 | 21.66 | 0.00 |
| | λ = 0 + Lat | 10 | 2718.49 | 230.08 | 0.00 |
| 17. Genome size × Habitat × Time | λ = 1 + Lat | 14 | 2504.72 | 16.30 | 0.00 |
| | λ = 0 + Lat | 14 | 2669.50 | 181.08 | 0.00 |
| 18. Body mass × Genome size | λ = 1 + Lat + Time | 7 | 2508.13 | 19.72 | 0.00 |
| | λ = 0 + Lat + Time | 7 | 2624.80 | 136.38 | 0.00 |
| | λ = 1 + Lat | 6 | 2524.21 | 35.80 | 0.00 |
| | λ = 1 + Time | 6 | 2548.68 | 60.27 | 0.00 |
| 19. Body mass × Genome size × Breathing mode | λ = 1 + Lat + Time | 11 | 2507.55 | 19.13 | 0.00 |
| | λ = 0 + Lat + Time | 11 | 2570.19 | 81.78 | 0.00 |
| | λ = 1 + Lat | 10 | 2523.09 | 34.68 | 0.00 |
| | λ = 1 + Time | 10 | 2536.95 | 48.54 | 0.00 |
| 20. Body mass × Genome size × Life-stage | λ = 1 + Lat + Time | 11 | 2505.94 | 17.53 | 0.00 |
| | λ = 0 + Lat + Time | 11 | 2620.59 | 132.18 | 0.00 |
| | λ = 1 + Lat | 10 | 2517.28 | 28.86 | 0.00 |
| | λ = 1 + Time | 10 | 2544.61 | 56.20 | 0.00 |
| 21. Body mass × Genome size × Habitat | λ = 1 + Lat + Time | 15 | 2517.56 | 29.15 | 0.00 |
| | λ = 0 + Lat + Time | 15 | 2583.76 | 95.34 | 0.00 |
| | λ = 1 + Lat | 14 | 2534.34 | 45.92 | 0.00 |
| | λ = 1 + Time | 14 | 2548.99 | 60.58 | 0.00 |
| **22. Body mass × Genome size × Breathing mode × Time** | **λ = 1 + Lat** | **18** | **2490.36** | **1.95** | **0.22** |
| | λ = 0 + Lat | 18 | 2538.30 | 49.89 | 0.00 |
| 23. Body mass × Genome size × Life-stage × Time | λ = 1 + Lat | 18 | 2501.61 | 13.20 | 0.00 |
| | λ = 0 + Lat | 18 | 2623.35 | 134.94 | 0.00 |

**Table S8.** Summary outputs for phylogenetic generalized least squares (PGLSs) models with the highest support ($w_i$) for CTmax analysis enlisted in Table S7. Models considered as covariate the absolute latitude (Lat) of animal collection. Pagel's lambda $\lambda = 1$ denotes correlation structure used (Brownian phylogeny).

| **Model** | **Variable** | **Estimate** | **SE** | ***P*-value** |
|---|---|---|---|---|
| Body mass × Breathing mode × Time + Lat ($\lambda$ = 1); AICc = 2488.41; $w_i$ = 0.57; ΔAICc = 1.00 | Intercept | 42.26 | 13.59 | 0.0020 |
| | **Body mass** | **-2.88** | **0.87** | **0.0011** |
| | Water | -0.20 | 2.24 | 0.9272 |
| | Time | 0.75 | 1.16 | 0.5177 |
| | **Lat** | **-0.08** | **0.01** | **0.0000** |
| | **Body mass × Water** | **4.99** | **1.07** | **0.0000** |
| | **Body mass × Time** | **1.43** | **0.47** | **0.0027** |
| | Water × Time | -1.51 | 1.19 | 0.2072 |
| | **Body mass × Water × Time** | **-2.72** | **0.58** | **0.0000** |
| Body mass × Genome size × Breathing mode × Time ($\lambda$ = 1); AICc = 2490.36; $w_i$ = 0.22; ΔAICc = 1.95 | Intercept | 33.93 | 13.94 | 0.0156 |
| | Body mass | -3.13 | 1.75 | 0.0737 |
| | Genome size | 18.79 | 11.00 | 0.082 |
| | **Water** | **7.93** | **4.25** | **0.0623** |
| | **Time** | **5.71** | **2.32** | **0.0143** |
| | **Lat** | **-0.08** | **0.01** | **0.0000** |
| | **Body mass × Genome size** | **11.35** | **4.13** | **0.0062** |
| | **Body mass × Water** | **5.34** | **1.89** | **0.0050** |
| | Genome size × Water | -17.56 | 11.09 | 0.1143 |
| | Body mass × Time | 1.13 | 1.06 | 0.2880 |
| | Genome size × Time | -11.81 | 6.96 | 0.0905 |
| | **Water × Time** | **-6.45** | **2.34** | **0.0061** |
| | **Body mass × Genome size × Water** | **-12.54** | **4.61** | **0.0068** |
| | **Body mass × Genome size × Time** | **-7.90** | **2.57** | **0.0023** |
| Body mass × Life-stage × Time + Lat ($\lambda$ = 1); AICc = 2490.42; $w_i$ = 0.21; ΔAICc = 2.00 | Intercept | 43.66 | 13.48 | 0.0013 |
| | Body mass | -0.95 | 0.57 | 0.0957 |
| | Non-adult | -1.42 | 1.39 | 0.3057 |
| | **Time** | **-1.20** | **0.31** | **0.0001** |
| | **Lat** | **-0.10** | **0.01** | **0.0000** |
| | **Body mass × Non-adult** | **5.26** | **1.13** | **0.0000** |
| | Body mass × Time | 0.34 | 0.27 | 0.2161 |
| | Non-adult × Time | 1.32 | 0.85 | 0.1197 |
| | **Body mass × Non-adult × Time** | **2.69** | **0.66** | **0.0001** |

**Table S9.** Selection results for phylogenetic generalized least squares (PGLSs) models to explain variation in ectotherms' CTmin (N = 190 species) as function of $log_{10}$–transformed body mass, $log_{10}$–transformed genome, breathing mode (air and water), life-stage (adults and non-adults), habitat (aquatic, intertidal and terrestrial) and all their possible interactions. Each model was assessed using as covariates the exposure duration (Time) and/or absolute latitude (Lat) of animal collection. Pagel's lambda (λ) denotes correlation structure used (λ = 0, star phylogeny and λ = 1, Brownian phylogeny). Number of parameters (*k*), corrected Akaike's information criterion (AICc), difference in AICc respect to the model with highest support (ΔAICc) and the Akaike's weights ($w_i$) are mentioned for each model.

| **Models** | | ***k*** | **AICc** | **ΔAICc** | $w_i$ |
|---|---|---|---|---|---|
| 0. Covariates | λ = 1 + Lat | 3 | 1076.21 | 87.32 | 0.00 |
| | λ = 0 + Lat | 3 | 1157.44 | 168.55 | 0.00 |
| 1. Body mass | λ = 1 + Lat | 4 | 1078.03 | 89.14 | 0.00 |
| | λ = 0 + Lat | 4 | 1050.47 | 61.58 | 0.00 |
| 2. Breathing Mode | λ = 1 + Lat | 4 | 1076.83 | 87.94 | 0.00 |
| | λ = 0 + Lat | 4 | 1150.57 | 161.68 | 0.00 |
| 3. Life-stage | λ = 1 + Lat | 4 | 1077.85 | 88.95 | 0.00 |
| | λ = 0 + Lat | 4 | 1158.68 | 169.79 | 0.00 |
| 4. Habitat | λ = 1 + Lat | 5 | 1063.79 | 74.90 | 0.00 |
| | λ = 0 + Lat | 5 | 1148.50 | 159.61 | 0.00 |
| 5. Body mass × Breathing Mode | λ = 1 + Lat | 6 | 1074.25 | 85.35 | 0.00 |
| | λ = 0 + Lat | 6 | 1029.96 | 41.06 | 0.00 |
| 6. Body mass × Life-stage | λ = 1 + Lat | 6 | 1076.77 | 87.88 | 0.00 |
| | λ = 0 + Lat | 6 | 1045.05 | 56.16 | 0.00 |
| 7. Body mass × Habitat | λ = 1 + Lat | 8 | 1069.34 | 80.45 | 0.00 |
| | λ = 0 + Lat | 8 | 1043.80 | 54.91 | 0.00 |
| 8. Body mass × Breathing mode × Time | λ = 1 + Lat | 10 | 1051.86 | 62.97 | 0.00 |
| | λ = 0 + Lat | 10 | 1011.38 | 22.49 | 0.00 |
| 9. Body mass × Breathing mode × Time | λ = 1 + Lat | 10 | 1052.20 | 63.30 | 0.00 |
| | λ = 0 + Lat | 10 | 1019.62 | 30.73 | 0.00 |
| 10. Body mass × Breathing mode × Time | λ = 1 + Lat | 14 | 1039.66 | 50.77 | 0.00 |
| | λ = 0 + Lat | 14 | 1039.08 | 50.19 | 0.00 |
| 11. Genome size | λ = 1 + Lat | 4 | 1074.03 | 85.13 | 0.00 |
| | λ = 0 + Lat | 4 | 1159.53 | 170.63 | 0.00 |
| 12. Genome size × Breathing mode | λ = 1 + Lat | 6 | 1070.58 | 81.68 | 0.00 |
| | λ = 0 + Lat | 6 | 1144.42 | 155.53 | 0.00 |
| 13. Genome size × Life-stage | λ = 1 + Lat | 6 | 1075.65 | 86.76 | 0.00 |
| | λ = 0 + Lat | 6 | 1154.62 | 165.72 | 0.00 |
| 14. Genome size × Habitat | λ = 1 + Lat | 8 | 1046.08 | 57.19 | 0.00 |
| | λ = 0 + Lat | 8 | 1153.09 | 164.19 | 0.00 |
| 15. Genome size × Breathing mode × Time | λ = 1 + Lat | 10 | 1047.97 | 59.08 | 0.00 |
| | λ = 0 + Lat | 10 | 1136.59 | 147.70 | 0.00 |
| 16. Genome size × Life-stage × Time | λ = 1 + Lat | 10 | 1045.99 | 57.10 | 0.00 |
| | λ = 0 + Lat | 10 | 1156.36 | 167.47 | 0.00 |
| 17. Genome size × Habitat × Time | λ = 1 + Lat | 14 | 1038.56 | 49.67 | 0.00 |
| | λ = 0 + Lat | 14 | 1153.79 | 164.89 | 0.00 |
| 18. Body mass × Genome size | λ = 1 + Lat | 6 | 1063.69 | 74.80 | 0.00 |
| | λ = 0 + Lat | 6 | 1046.91 | 58.02 | 0.00 |
| **19. Body mass × Genome size × Breathing mode** | λ = 1 + Lat | 10 | 1060.58 | 71.69 | 0.00 |
| | **λ = 0 + Lat** | **10** | **988.89** | **0.00** | **0.99** |
| 20. Body mass × Genome size × Life-stage | λ = 1 + Lat | 10 | 1054.80 | 65.91 | 0.00 |
| | λ = 0 + Lat | 10 | 1015.53 | 26.63 | 0.00 |
| 21. Body mass × Genome size × Habitat | λ = 1 + Lat | 14 | 1050.25 | 61.36 | 0.00 |
| | λ = 0 + Lat | 14 | 1045.22 | 56.33 | 0.00 |

**Table S9 continued**

| Models | | $k$ | AICc | ΔAICc | $w_i$ |
|---|---|---|---|---|---|
| 22. Body mass × Genome size × Breathing mode × Time | λ = 1 + Lat | 18 | 1048.76 | 59.87 | 0.00 |
| | λ = 0 + Lat | 18 | 997.91 | 9.02 | 0.01 |
| 23. Body mass × Genome size × Life-stage × Time | λ = 1 + Lat | 18 | 1040.60 | 51.71 | 0.00 |
| | λ = 0 + Lat | 18 | 1019.33 | 30.44 | 0.00 |

**Table S10.** Summary outputs for the phylogenetic generalized least squares (PGLSs) model with the highest support ($w_i$) for CTmin analysis enlisted in Table S9. Model considered as covariate the absolute latitude (Lat) of animal collection. Pagel's lambda $\lambda = 0$ denotes correlation structure used (star phylogeny).

| Model | Variable | Estimate | SE | ***P*-value** |
|---|---|---|---|---|
| Body mass × Genome size × Breathing mode + Lat ($\lambda = 0$) | Intercept | 9.64 | 0.92 | 0.0000 |
| | **Body mass** | **3.21** | **0.37** | **0.0000** |
| | Genome size | -2.52 | 2.13 | 0.2385 |
| | **Water** | **3.66** | **0.90** | **0.0001** |
| | **Lat** | **-0.22** | **0.02** | **0.0000** |
| | **Body mass × Genome size** | **4.26** | **1.13** | **0.0002** |
| | **Body mass × Water** | **-3.33** | **0.71** | **0.0000** |
| | Genome size × Water | 0.00 | 2.47 | 0.9988 |
| | Body mass × Genome size × Water | -0.73 | 1.86 | 0.6945 |

**Table S11.** Values of phylogenetic signal, estimated as Pagel's lambda (λ), for continuous traits used in the models' correlation structure. For each variable, the log-likelihood (*LL*) is mentioned. *P*-values correspond to the significance of the observed λ compared to one expected under Brownian motion (λ = 1).

| **Variable** | **Number of species** | **Pagel's lambda (λ)** | ***LL*** | **P-value** |
|---|---|---|---|---|
| CTmax | 510 | 0.92 | -1504.77 | $6.75 \times 10^{-76}$ |
| CTmin | 232 | 0.96 | -698.88 | $1.35 \times 10^{-35}$ |
| Body mass | 510 | 0.89 | -722.12 | $2.85 \times 10^{-87}$ |
| Genome size | 433 | 0.92 | -77.31 | $3.64 \times 10^{-61}$ |
| Time (for CTmax) | 510 | 0.79 | -341.47 | $1.73 \times 10^{-33}$ |
| Time (for CTmin) | 232 | 0.90 | -113.43 | $3.18 \times 10^{-13}$ |
| Lat | 510 | 0.98 | -1958.15 | $1.62\times 10^{-87}$ |
